# A comprehensive portrait of antimicrobial resistance in the zoonotic pathogen *Streptococcus suis*

**DOI:** 10.1101/2020.05.05.078493

**Authors:** Nazreen F. Hadjirin, Eric L. Miller, Gemma G. R. Murray, Phung L. K. Yen, Ho D. Phuc, Thomas M. Wileman, Juan Hernandez-Garcia, Susanna M. Williamson, Julian Parkhill, Duncan J. Maskell, Rui Zhou, Nahuel Fittipaldi, Marcelo Gottschalk, A.W. (Dan) Tucker, Ngo Thi Hoa, John J. Welch, Lucy A. Weinert

## Abstract

Antimicrobial resistance (AMR) is among the gravest threats to human health and food security worldwide. Pigs receive more antimicrobials than most other livestock, and are a known source of zoonotic disease. We studied AMR in *Streptococcus suis*, a commensal found in most pigs, but which can also cause serious disease in both pigs and humans. We obtained replicated measures of Minimum Inhibitory Concentration (MIC) for 16 antibiotics, across a panel of 678 isolates, from the major pig-producing regions of the world. For several drugs, there was no natural separation into “resistant” and “susceptible”, highlighting the need to treat MIC as a quantitative trait. We found differences in MICs between countries, consistent with their patterns of antimicrobial usage. AMR levels were high even for drugs not used to treat *S. suis*, with many multi-drug resistant isolates. And similar levels of resistance were found in pigs and humans from zoonotic regions. We next used whole genome sequences for each isolate to identify 43 candidate resistance determinants, 22 of which were novel in *S. suis*. The presence of these determinants explained most of the variation in MIC. But there were also complications, including epistatic interactions, where known resistance alleles had no effect in some genetic backgrounds. Beta-lactam resistance involved many variants of small effect, appearing in a characteristic order. Our results confirm the potential for genomic data to aid in the fight against AMR, but also demonstrate that it cannot be tackled one species or one drug at a time.

## Introduction

The ability of bacterial pathogens to evolve resistance to antimicrobials is one of the gravest threats to human health and food security worldwide. Antimicrobial resistance (AMR) to a given drug can be quantified via the minimum inhibitory concentration (MIC), i.e., the concentration of the drug that is sufficient to inhibit growth of a bacterial culture. While, for practical use, bacterial isolates are often categorised as either susceptible or resistant, MIC is a continuously varying trait. In this study, we investigate how AMR genes, variants and ecology explain antibiotic phenotype in the bacterium *Streptococcus suis*, treating MIC phenotype as a quantitative trait.

*S. suis* primarily exists as a commensal in pigs, colonising the nasopharynx, gut and vagina, but it also causes systemic and respiratory disease, particularly in young pigs [1]. *S. suis* is also a serious zoonotic disease, being the leading cause of adult bacterial meningitis in Vietnam [2]. While some autogenous vaccines are used in pig production, they are serotype-specific and give inconsistent cross protection against heterogeneous *S. suis* [3]. Antimicrobials therefore remain the standard treatment for *S. suis*, and as such, *S. suis* is a leading driver of antimicrobial usage in pig farms [4].

As well as being a serious problem in itself, *S. suis* also has unique benefits as a model for studying AMR. By weight, more pork is consumed globally than any other meat [5, 6], and *S. suis* is found in most, if not all pigs [7]. Furthermore, antibiotic consumption is higher in pigs (172 mg per population corrected unit) than any other livestock (e.g. cattle (45 mg) and chicken (148 mg)) [8]. As a result, most *S. suis* lineages will experience antibiotics. These include not only antibiotics administered directly against *S. suis*, whether as a therapeutic, prophylaxis or metaphylaxis, but also, and perhaps more commonly, in response to many other bacterial infections.

The strong selection pressure caused by widespread use of antimicrobials in pig farming is expected to give rise to AMR in *S. suis*. Consistent with this, several phenotypic studies show high MICs for each of the major classes of antibiotics in one or more *S. suis* collections [9–13]. In addition, there have been demonstrations of individual resistance determinants affecting MIC in *S. suis* [14–16], and also some mining of *S. suis* genome collections for known resistance determinants [17–19]. However, to our knowledge, no study has combined complete genomic and phenotypic information in large numbers of isolates, collected from a diverse range of populations.

To this end, we obtained replicated measures of MIC for 16 antibiotics, each widely used in pigs, for six different *S. suis* collections, comprising 678 isolates. These collections were chosen to include the three main pig-producing regions of the globe - namely the Americas, South East Asia and Europe - and targeted both human and pig hosts, as well as a range of years, serotypes and clinical phenotypes [20–22]. We obtained whole genome sequences for each of our isolates, allowing us to compare phenotype and genotype on an unprecedented scale.

## Results

### 1. Measurement of MIC for 16 antibiotics in *S. suis*

We scored 678 isolates of *Streptococcus suis* for replicated measures of Minimum Inhibitory Concentration (MIC) for 16 antibiotics (Table S1). Four of these are beta-lactams (amoxicillin, cefquinome, ceftiofur and penicillin), which are typically used to treat *S. suis* infection in pigs. The other antibiotics are all widely used in the pig industry, but found in different drug classes. They comprise macrolide-lincosamide-streptogramin B (MLS_B_: erythromycin, lincomycin, tilmicosin and tylosin), tetracyclines (doxycycline and tetracycline), and fluoroquinolones (enrofloxacin and marbofloxacin), plus one each of an aminoglycoside (spectinomycin), a pleuromutilin (tiamulin), trimethoprim (TMP), and a phenicol (florfenicol). Results, shown in the left-hand plot of Figure 1, reveal wide variation among our isolates in MIC for most of the antibiotics. For a few antibiotics, especially the MLS_B_ and tetracyclines, the MIC values are clearly bimodal, suggesting a meaningful division between “resistant” and “wild-type” isolates. However, for most antibiotics, the distributions are either roughly lognormal (fluoroquinolones and phenicols) or positively skewed (spectinomycin, pleuromutilin, TMP and beta-lactams). It is striking that the distributions for the beta-lactams are not clearly dissimilar from any of the other antibiotic classes, despite their being the most common treatment against *S. suis*. Given these distributions, and the general lack of defined clinical breakpoints for these antibiotics in *S. suis* [23], we treated MIC as a quantitative trait in our subsequent analyses.

**Figure 1.**
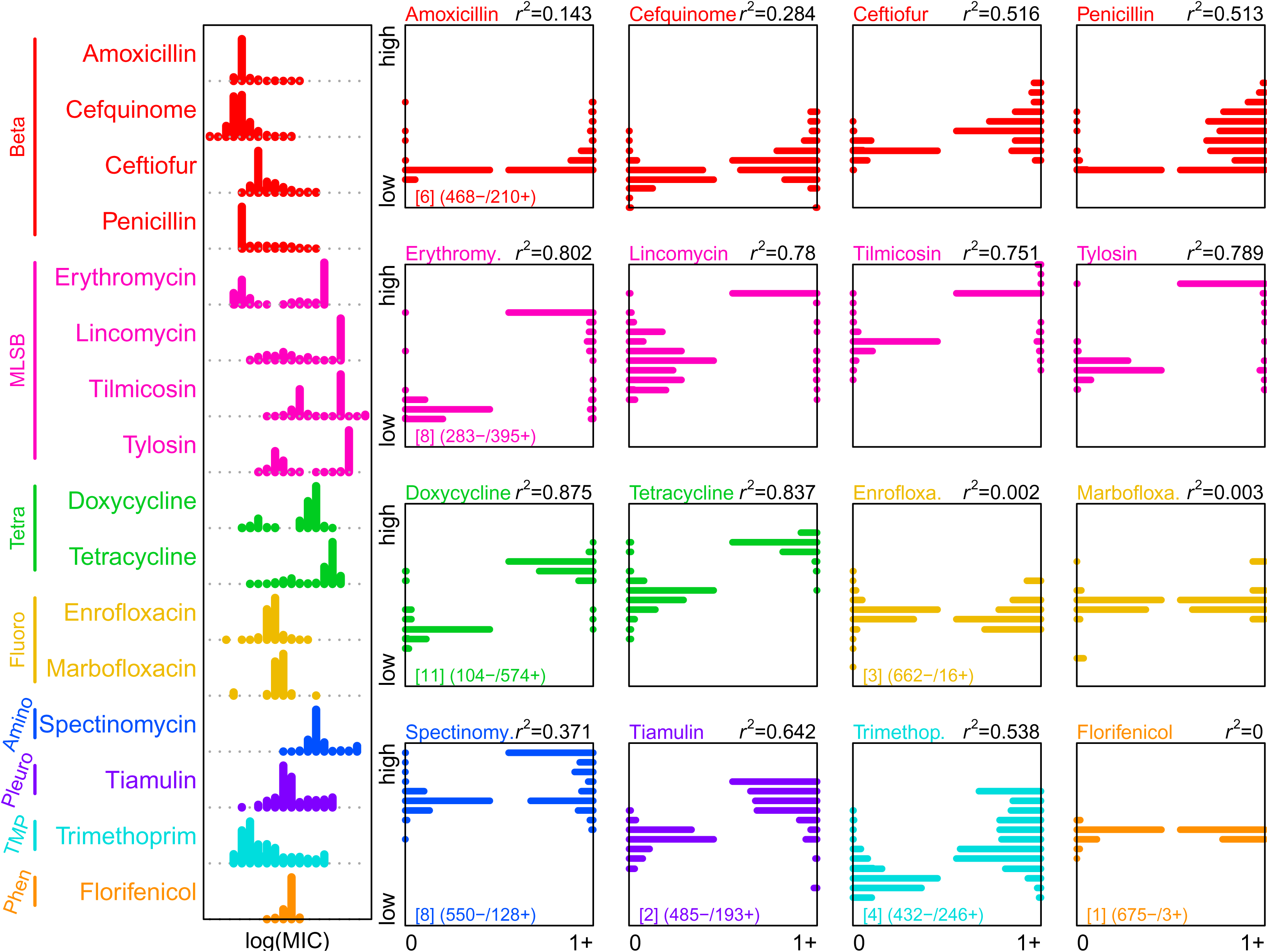
Candidate AMR determinants explain most of the variation in MIC. Histograms of log transformed MIC measures for each of our 16 different antibiotics, across our panel of 678 *S. suis* isolates. Antibiotics are coloured by their class (Beta: beta-lactams; MLS_B_: macrolide-lincosamide-streptogramin B; Tetra: tetracyclines; Fluoro: fluoroquinolones; Amino: aminoglycoside; Pleuro: pleuromutilin; TMP: trimethoprim and Phen: phenicol). In the 16 square panels, the left-hand histograms (labelled 0) show the MIC values for isolates that carry no determinant for that antibiotic class, while the right-hand histograms (labelled 1+) show the MIC values for isolates carrying one or more such determinant. If all resistance determinants perfectly explain MIC, then we expect to see histogram distributions on the bottom left (low MIC, no determinant) and the top right (high MIC, presence of candidate determinant(s)). For the first antimicrobial in each class, we show the number of candidate AMR determinants in square brackets, along with the number of isolates where candidate determinants are absent or present. *r*^2^ values show the proportion of the variance explained in a standard ANOVA by the presence of one or more candidate determinant.

### 2. Ecological and genomic predictors of MIC

Our *S. suis* isolates were collected at different times, and from different countries, hosts and body sites, including sites associated with respiratory and systemic disease. Isolates were also genetically heterogeneous, and differed in their serotype; including serotypes with a known disease association (see methods and Table S1). We first used linear models, to ask whether MIC levels varied systematically with these ecological and genomic factors.

*S. suis* is highly recombining, such that no single genealogy describes its diversity [20]. As such, to characterise genomic diversity in our sample, we identified 30 clusters of similar strains (see methods). The resulting clusters were a highly significant predictor of MIC (*p*<10^-15^; Table 1a) showing that genomically similar isolates tend to have similar MIC values. To account for this genetic structure, “cluster” was included as a random effect in subsequent analyses.

**Table 1.** Ecological and genomic predictors of MIC.

| Analysis | #strains | Fixed effects | Df | F | p |
| --- | --- | --- | --- | --- | --- |
| (a) | 678 | Antibiotic | 15 | 1404.125 | <10<sup>-15</sup> |
|  |  | genetic cluster | 29 | 40.644 | <10<sup>-15</sup> |
| (b) | 450 | Antibiotic | 15 | 972.3107 | <10<sup>-4</sup> |
|  |  | Year | 1 | 6.6078 | 0.0102 |
|  |  | Serotype | 1 | 6.6261 | 0.0101 |
|  |  | Disease Status | 2 | 2.9760 | 0.0511 |
|  |  | Country | 3 | 92.5155 | <10<sup>-4</sup> |
| (c) | 678 | Antibiotic | 15 | 1457.4043 | <10<sup>-4</sup> |
|  |  | Country | 3 | 139.5902 | <10<sup>-4</sup> |
| (d) | 557 | Antibiotic | 15 | 1144.3846 | <10<sup>-4</sup> |
|  |  | Disease Status | 2 | 9.9619 | <10<sup>-4</sup> |
| (e) | 542 | Antibiotic | 15 | 1131.6347 | <10<sup>-4</sup> |
|  |  | Serotype | 1 | 8.7216 | 0.0032 |
| (f) | 652 | Antibiotic | 15 | 1350.8781 | <10<sup>-4</sup> |
|  |  | Year | 1 | 0.1591 | 0.69 |

A model including all potential predictors showed that country of isolation was also a highly significant predictor of MIC. In addition, there were weaker effects of year of collection (with a slight trend for increasing MIC over time); for serotype (with non-disease-associated serotypes tending to have higher MIC); and for host disease status (with non-clinical isolates having higher MIC) (Table 1b).

Separate analyses for each antibiotic, showed that the effect of country was driven by consistently higher MIC in the samples from Canadian pigs. As shown in Figure 2a (left-hand panel), this applied to 15/16 antibiotics, and also applied to various models (Table 1c), and subsets of the data (Figure S1). For example, the effect is seen consistently in systemic pathogens, respiratory pathogens and non-clinical isolates (Figure S1). By contrast, the other predictors (year, serotype and clinical status) had effects that were less consistent or robust (Table 1d-f). The difference in the MIC values between Canada and the UK is consistent with higher antimicrobial usage in Canada [24, 25].

**Figure 2.**
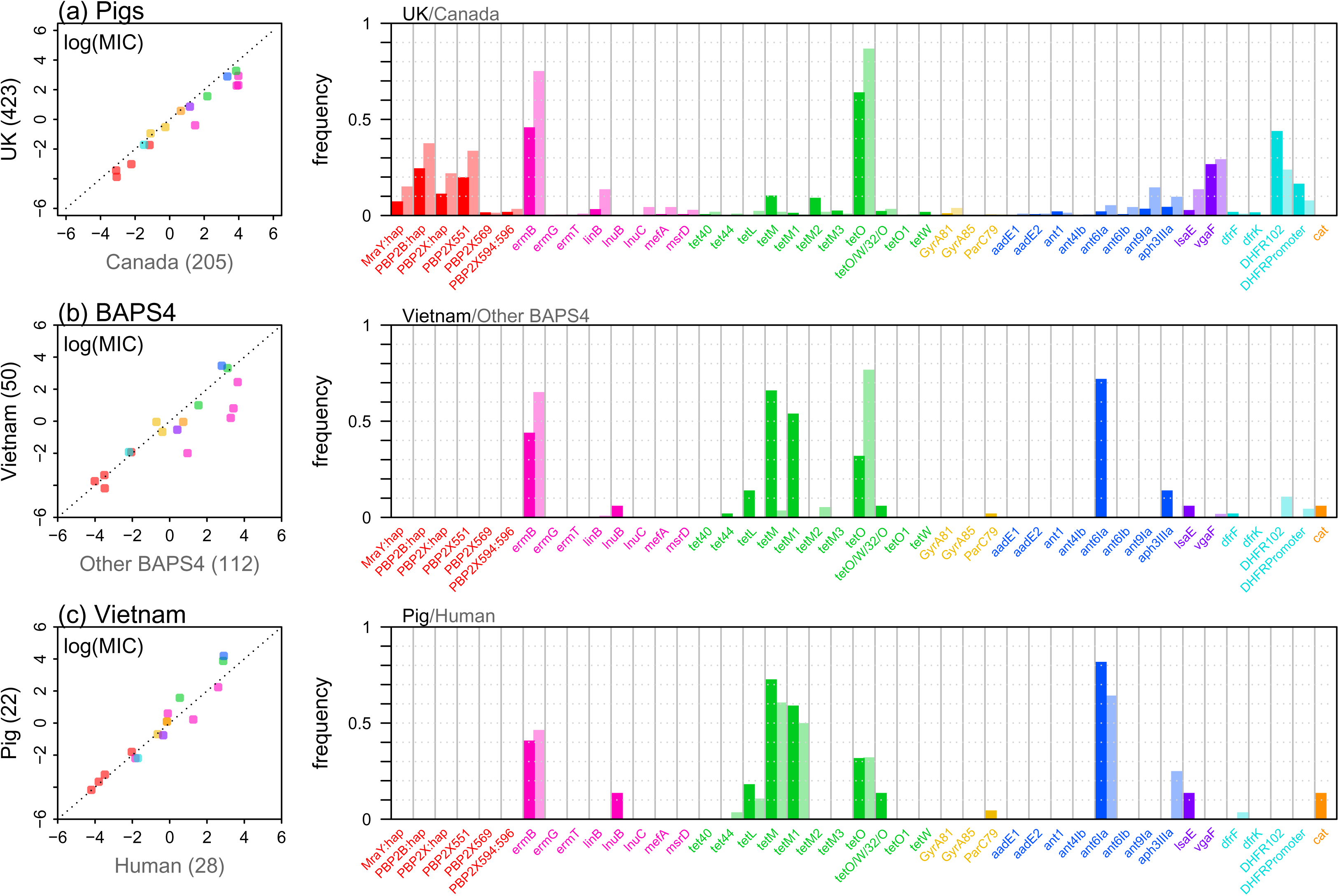
Differences in MICs between subsets of the data. Each row compares two subsets of the isolates: (a) the 423 isolates from UK pigs, and the 205 isolates from Canadian pigs. (b) the 50 isolates from Vietnam (all of which are from the genetic cluster “BAPS4”), and the 112 BAPS4 isolates from the UK and Canada. (c) the 22 Vietnamese isolates from pigs, and the 28 Vietnamese isolates from people. Left-hand panels compare the mean log MICs for each antibiotic. Consistent deviations from the dotted 1:1 line suggest consistently higher or lower MICs in that subset of the data (so the rightward shift in panel (a) shows that MICs are consistently higher in Canada). Right-hand panels show the proportion of isolates that carry each of the 43 candidate AMR determinants. Each point or bar is coloured according to its drug class, according to the colour scheme in Figure 1.

Predictions for Vietnam are more difficult, as there is a lack of official records of antibiotic usage, although it is likely that usage is both higher than in the EU and focussed on different drugs [26]. However, as shown in Figure 2b (left-hand panel), we saw no such signal in the MIC data, even when we restricted the comparison to the genetic cluster BAPS4 that includes all of our Vietnamese sample (see methods). The clearest pattern is for lower MIC for the MLS_B_ class in the Vietnamese isolates (pink points in Figure 2b left-hand panel). As shown in Figure 2c (left-hand panel), we also found no robust differences between Vietnamese isolates sampled from humans and pigs. This is consistent with pigs being a reservoir for this zoonotic disease, and the lack of evidence of genomic adaptation to human hosts [20].

### 3. Identification of candidate AMR determinants within *S. suis* genomes

Using a candidate locus approach, we next identified known and putative determinants of AMR that were present in our sampled genomes. We considered both the presence/absence of whole genes, and sequence variants within genes. In total, we detected 43 determinants: 10 completely novel variants of known resistance genes, 12 known resistance genes detected in *S. suis* for the first time, and 21 previously reported in *S. suis* (Table 2 and Table S1). Our candidates come from three sources: the antimicrobial resistance database CARD [27], previously published *S. suis* variants not present in the CARD database [28–30] and novel variants detected using a subset of our data (see methods).

**Table 2.** AMR determinants of Streptococcus suis.

| AMR gene | Name in figures | Resistance conferred (Antibiotic Class) | No. of isolates | Reported in *S. suis* | Reported in other species (CARD and NCBI) | Reported genetic location in *S. suis* or other species |
| --- | --- | --- | --- | --- | --- | --- |
| *ant(4')-Ib* | ant4Ib | Aminoglycoside | 1 | This report | *Staphylococcus aureus*, *Staphylococcus epidermidis*, *Enterococcus spp.* | Plasmid |
| *ant(6')-Ia/aadE* | ant6Ia | Aminoglycoside | 56 | [19] | *Staphylococcus epidermidis*, *Streptococcus spp.*, *Enterococcus faecalis* | Plasmid, ICE |
| *ant(6')-Ib* | ant6Ib | Aminoglycoside | 12 | This report | *Campylobacter spp.*, *Bacillus subtilis*, *Clostridium spp.* | Pathogenicity Island |
| *ant(9')-Ia/spw/aad9* | ant9Ia | Aminoglycoside | 45 | [19] | *Staphylococcus aureus*, *Enterococcus spp.*, *Staphylococcus spp.* | Plasmid, ICE |
| *aph(3')-IIIa* | aph3IIIa | Aminoglycoside | 46 | [85] | *Campylobacter spp.*, *Staphylococcus spp.*, *Streptococcus spp.*, *Enterococcus spp.* | Plasmid, transposon |
| *ant1\** | ant1 | Aminoglycoside | 12 | This report | *Campylobacter spp.*, *Clostridium difficile*, *Lactobacillus amylovorus* | Plasmid, ICE |
| *aadE1\** | aadE1 | Aminoglycoside | 2 | This report | *Enterococcus faecium*, *Staphylococcus aureus* | Plasmid |
| *aadE2\** | aadE2 | Aminoglycoside | 5 | This report | *Clostridium spp.*, *Campylobacter fetus* | ICE |
| *ermB* | ermB | MLS<sub>B</sub> (Macrolide/<br>Lincosamide) | 370 | [86] | *Campylobacter spp.*, *Clostridium spp.*, *Enterococcus spp.*, *Escherichia coli*, *Klebsiella pneumoniae*, *Proteus mirabilis*, *Shigella spp.*, *Staphylococcus spp.*, *Streptococcus spp.*, *Vibrio vulnificus*. | Plasmid, ICE |
| *ermG* | ermG | MLS<sub>B</sub> (Macrolide/<br>Lincosamide) | 1 | This report | *Clostridium difficile*, *Enterococcus faecium* | Conjugative transposon |
| *ermT* | ermT | MLS<sub>B</sub> (Macrolide/<br>Lincosamide) | 3 | This report | *Enterococcus faecium*, *Staphylococcus aureus*, *Streptococcus spp.* | Plasmid |
| *mefA* | mefA | Macrolide | 11 | [86] | *Streptococcus spp.*, *Enterococcus faecium*, *Haemophilus parainfluenzae*, *Proteus mirabilis* | Transposon, ICE |
| *mel/msrD* | mel | Macrolide | 9 | [86] | *Streptococcus spp.*, *Enterococcus faecium*, *Haemophilus parainfluenzae*, *Proteus mirabilis*. | Transposon, ICE |
| *lnuB* | lnuB | Lincosamide | 3 | [17] | *Enterococcus spp.*, *Staphylococcus spp.*, *Streptococcus agalactiae* | Plasmid |
| *linB* | linB | Lincosamide | 42 | [87] |  |  |
| *lnuC* | lnuC | Lincosamide | 9 | [88] | *Campylobacter spp.*, *Clostridium difficile*, *Streptococcus agalactiae*. | Plasmid |
| *lsaE* | lsaE | Pleuromutilin | 43 | [17] | *Enterococcus spp.*, *Staphylococcus spp.*, *Streptococcus agalactiae*. | Plasmid, ICE |
| *tet44* | tet44 | Tetracycline | 4 | This report | *Clostridium spp.* | ICE |
| *tetM* | tetM | Tetracycline | 81 | [19] | *Streptococcus spp.*, *Escherichia coli*, *Staphylococcus spp.*, *Enterococcus spp.*, *Klebsiella pneumoniae*, *Neisseria meningitidis*, *Clostridium difficile*, *Salmonella enterica* | Plasmid, ICE |
| *tetM1\** | tetM1 | Tetracycline | 34 | This report | *Staphylococcus spp.*, *Streptococcus spp.*, *Enterococcus spp.*, *Bacillus spp.*, *Clostridium spp.* | Plasmid, transposon |
| *tetM2\** | tetM2 | Tetracycline | 43 | This report | *Enterococcus spp.*, *Clostridium difficile*. |  |
| *tetM3\** | tetM3 | Tetracycline | 13 | This report | *Streptococcus pneumoniae*, *Streptococcus pyogenes*, *Enterococcus spp.*, *Clostridium spp.* | Transposon, ICE |
| *tetO* | tetO | Tetracycline | 465 | [19] | *Campylobacter spp.* | ICE, plasmid |
| *tetO1\** | tetO1 | Tetracycline | 2 | This report | *Dorea spp.*, uncultured bacterium | Conjugative transposon |
| *tetW* | tetW | Tetracycline | 8 | [16] | *Streptococcus agalactiae*, *Clostridium difficile*, *Chlamydia trachomatis*. | Plasmid |
| *tet40* | tet40 | Tetracycline | 7 | [17, 29] | *Clostridium difficile* | Composite mobile genetic element, ICE |
| *tetL* | tetL | Tetracycline | 12 | [19] | *Streptococcus spp.*, *Escherichia coli*, *Staphylococcus spp.*, *Enterococcus spp.*, *Klebsiella pneumoniae*. | Plasmid |
| *tet(O/W/32/O) †* | tetO:W:32:O | Tetracycline | 20 | [84] |  | ICE |
| *dfrF* | dfrF | TMP | 11 | This report | *Clostridium difficile*, *Enterococcus spp.*, *Staphylococcus aureus*, *Streptococcus agalactiae* | plasmid |
| *dfrK* | dfrK | TMP | 7 | This report | *Staphylococcus aureus*, *Staphylococcus epidermidis*. | plasmid |
| *vgaF\** | vgaF | Pleuromutilin | 173 | This report | Not known | Chromosome. Found as an intact gene (~1386bp) or truncated (<750bp) |
| *cat\** | cat | Florfenicol | 3 | This report | *Enterococcus faecalis*, *Streptococcus iniae*, *Listeria monocytogenes*, *Staphylococcus aureus* | Plasmid, Transposon |
| *dhfr\** | DHFR102 | TMP | 235 | This report | Mutations in I100L in *Streptococcus spp.* and *Staphylococcus aureus* | DHFR: I102L |
| *dhfr promoter\** | DHFRPromoter | TMP | 86 | This report | None reported | DHFR : A5G upstream/indels 0-30bp upstream |
| *pbp2B/penA\** | PBP2B:hap | Penicillin | 181 | This report | Mutations in other locations in *Streptococcus pneumonia* and Group B *Streptococcus spp* | PBP2B : K479T/A, D512E/Q/K/A, K513E/D,T515S |
| *pbp2X\** | PBP2X:hap | Ceftiofur | 93 | This report | None reported | PBP2X : M437L, S445T, T467S Y525F |
| *pbp2X* | PBP2X551 | Penicillin | 153 | This report | Mutation in PBP2X in *S. pneumoniae* (T550), group B streptococci (T555) and *S. pyogenes* (T553) | PBP2X : T551S |
| *pbp2X* | PBP2X594:596 | Penicillin | 15 | [28] | *Streptococcus suis*. Mutations in other locations in *Streptococcus pneumonia*, Group B *Streptococcus sp.* and *Streptococcus uberis* | PBP2X : L594Y/F, V596G |
| *pbp2X* | PBP2X569 | Penicillin | 10 | [28] |  | PBP2X : N569Q |
| *mraY\** | MraY:hap | Penicillin | 62 | This report | None reported | *MraY* : M6I/L and either A4S/T or G8S |
| *parC†* | ParC79 | Fluoroquinolone | 4 | [30] | *Escherichia coli*, *Pseudomonas aeruginosa*, *Streptococcus spp.*, *Staphylococcus aureus* | *ParC* : S79 |
| *gyrA†* | GyrA81 | Fluoroquinolone | 13 | [30] | *Escherichia coli*, *Klebsiella spp.*, *Streptococcus spp.*, *Staphylococcus aureus* | GyrA : S81 |
| *gyrA†* | GyrA85 | Fluoroquinolone | 1 | [30] | *Escherichia coli*, *Klebsiella spp.*, *Streptococcus spp.*, *Staphylococcus aureus* | GyrA : E85 |

\*novel determinants reported in this study † Not documented in the CARD database MLS<sub>B</sub> Macrolide Linocosome Streptogramin B TMP Trimethoprim

Of the 10 novel determinants, six are variants in genes previously associated with resistance in other bacteria (including a promoter variant). In particular, we discovered four novel haplotypes within the loop region of the central transpeptidase domains of the *pbp2B* and *pbp2X* genes and within the signal peptide region of the *mraY* gene (encoding the enzyme, acetylmuramoyl-pentapeptide-transferase, essential to cell wall biosynthesis) that were associated with variation in beta-lactam MIC (Table 2). In particular, we noted a PBP2X mutation at the conserved location T551, similar to that found in *S. pneumoniae* (T550), group B streptococci (T555) and *S. pyogenes* (T553) PBP2X proteins [31] indicative of a shared overall mechanism mediating beta-lactam resistance across the genus. Polymorphisms in *mraY* have also been identified in a genome-wide association study of beta-lactam resistance in *S. pneumoniae* [32]. In addition to these four beta-lactam haplotypes, two variants – a variant of the chromosomal dihydrofolate reductase gene *dhfr* and its promoter – were associated with reduced susceptibility against trimethoprim (Table 2). Furthermore, all of these six variants were independently associated with changes in MIC in at least seven different genetic clusters (Table S2) implying that the association between these variants and MIC is either directly causal or compensatory to the causal variant [33].

The remaining four novel variants were whole genes with homologies to known resistance genes: three located on mobile genetic elements (MGEs) and one chromosomal (Table 2). Three novel aminoglycoside genes, carried by isolates with high spectinomycin MIC, were identified based on aminoglycoside resistance determinant protein homologies. Although previously undescribed, we found homologues in other bacteria using a *blastn* search of the non-redundant database in *Genbank* (Table 2). We also characterised a chromosomally-encoded *vgaC* homologue (37% protein homology to *vgaC*), encoding an ABC-F ATP-binding cassette ribosomal protection protein, that we designate *vgaF.* This gene has arisen in 13 different BAPS clusters, each time associated with reduced susceptibility to tiamulin (Table S2).

The 12 previously-known AMR genes detected in *S. suis* for the first time are mobile genetic element (MGE)-linked genes that confer resistance to aminoglycosides (2/12), MLS_B_ (2/12), tetracyclines (5/12), TMP (2/12) and phenicol (1/12). Based on the CARD and NCBI databases, these genes are associated with other gram-positive and gram-negative bacteria (Table 2).

Overall, most of our candidate determinants (28/43) were found in fewer than 5% of the isolates, but *ermB* (which confers resistance to the MLS_B_ class) and *tetO* (which confers resistance to tetracyclines) were present in a majority of the isolates (Table 2). Only about a quarter of isolates carry a candidate determinant for the beta-lactams highlighting the continuing susceptibility to these first line treatment drugs in *S. suis*.

Nevertheless, as shown in Figure 3, many isolates carried multiple determinants, and multi-drug resistance was widespread. Around 40% of isolates carried resistance determinants to three or more classes of drug (275/678), and over 10% carried resistance determinants to five classes (81/678); this is more than carried no determinants at all (59/678).

**Figure 3.**
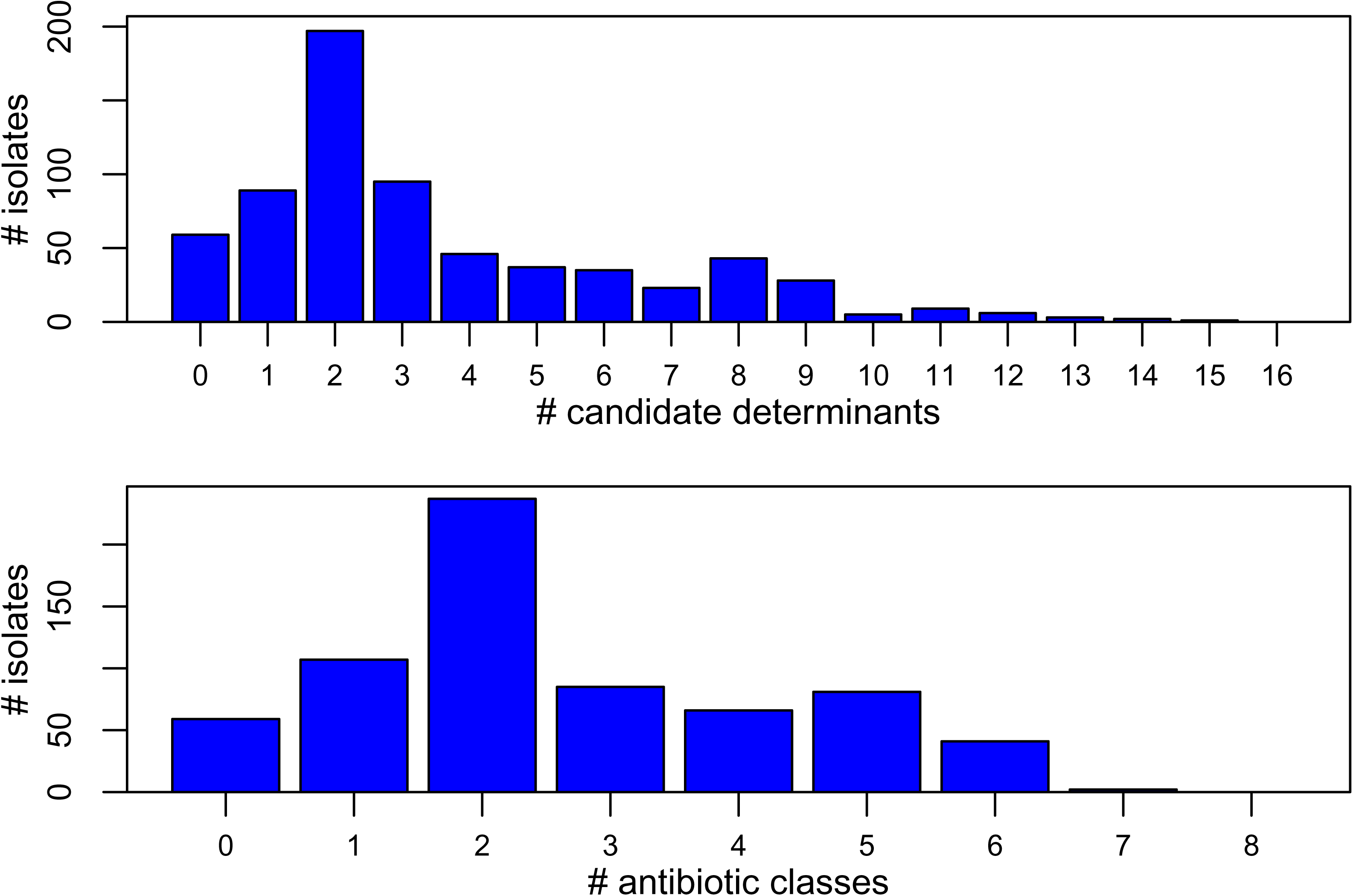
High levels of multidrug resistance in *S. suis*. The upper panel shows the number of our 678 isolates that carry a given number of candidate AMR determinants. The low panel shows the number of isolates that carry one or more AMR determinant for a given number of drug classes. Results show that more isolates carry determinants against 5 drug classes than carry no determinant at all.

### 4. Candidate AMR determinants explain most of the variation in MIC

We next investigated how well our candidate variants explain the variation we observe in MIC (see Figures S2-S8 for detailed plots). The right-hand panel of Figure 1 compares the distribution of MIC values for isolates carrying one or more of the candidate AMR determinants for that antibiotic class (denoted “1+”), to the MIC of the remaining isolates, which carry no such determinant (denoted “0”).

The proportion of the variation in MIC explained by our variants varied between antibiotics. Explanatory power was strongest (*r^2^*>0.75) for antibiotics where the distributions of MIC were most clearly bimodal (MLS_B_ and tetracyclines). By contrast, we found no association between our variants and MIC for the fluoroquinolones and phenicol, where the distribution of MIC values was roughly lognormal, consistent with our isolates representing a “wild-type” population. For the remaining antibiotics, our determinants had intermediate explanatory power (Figure 1), but there were very few isolates with high MIC that did not carry a candidate determinant. This suggests that we have detected most of the causal variants in our dataset, removing the need to perform additional genome-wide associations.

We next asked whether the variable presence of our candidate determinants in different ecological settings would explain the significant ecological predictors of MIC in our linear models (Table 1). Consistent with this hypothesis, the typical MIC level in each genetic cluster was highly correlated with the frequency of determinants in that cluster (Figure S9).

Differences between countries were explained in the same way. The higher MIC in Canada than the UK was due to consistently higher frequencies of the same determinants (Figure 2a right-hand panel) – with the exception of trimethoprim (see below). By contrast, tetracycline resistance throughout the BAPS4 cluster, though at similar levels in all three countries, was conferred by different determinants in Vietnam (Figure 2b).

### 5. Ineffective variants and epistasis

While our data contained few isolates with high MIC that did not carry a candidate determinant, there were many isolates with low MIC despite carrying a determinant (Figure 1).

In some cases, this was due to previously identified candidate genes that had no appreciable effect on MIC in our data. For example, spectinomycin MIC was unchanged by some determinants identified from the CARD database (e.g. *ant(6’)-Ib* and *aph(3’)-IIIa* in Figure S6). If we allow for the presence of these non-functional genes, the *r^2^* value for spectinomycin increases from 0.37 (Figure 1) to 0.69 (Figure S10a).

In other cases, as is well known, variants act against only some of the antibiotics in a class. In MLS_B_, for example, modification of the ribosomal target confers cross resistance to macrolides and lincosamides, while mechanisms such as efflux and enzymatic inactivation do not [34]; so *msrD* acts against erythromycin, but not against lincomycin, while *linB* inactivates lincomycin, but not erythromycin (Figure S3). Again, taking this into account further increases predictive power (Figure S10a).

As well as these simple cases, there were some clear examples of genetic interactions. For trimethoprim, two candidate variants – in the protein DHFR102 and its promoter (Table 2) – were found at higher frequency in the UK than Canada (Figure 2a right-hand-panel), but in Canada, the two variants were more often found together, leading to higher MIC overall (Figure 2a; Figure S8).

Finally, we observed complex patterns for the second most common determinant in our dataset, *ermB*. For the 334 isolates that carried only *ermB* (and no other candidate determinant to MLS_B_ class drugs), MIC values were clearly bimodal, with many carriers having very low MIC (Figures S3 and S11). Further analysis revealed a small number of isolates with frameshift or premature stop codons (4/326 isolates with complete sequences in our assemblies). While the remaining isolates with low MIC carried rare amino acid variants at one of four positions (T75X, N100S, R118H and V226I) – the last three of which differentiate the *ermB* sequences found in *Streptococcus pneumoniae* from *Clostridium perfringens* [35]. Nevertheless, no particular sequence was always associated with low MIC, suggesting some unidentified source of epistasis (see methods and Figure S11 for full details).

### 6. Resistance to beta-lactams

While beta-lactams are the major treatment class for *S. suis*, it is notable that the explanatory power of our variants seems to be weaker for this drug class than for, e.g. tetracyclines or MLS_B_.

In Streptococci a typical route to beta-lactam resistance involves variants in the *pbp* genes, and it is well established that the joint action of many *pbp* variants is necessary to explain substantial changes in MIC [36, 37]. Single point mutations in *pbp2x,* for example, cause very modest elevations in *S. pneumoniae* [36], group B Streptococci [38], *Streptococcus dysgalactiae subsp. equisimilis* [39] and *Streptococcus pyogenes* [31].

Like *S. pneumoniae*, *S. suis* has three key mosaic *pbp* genes, *pbp1A, pbp2b* and *pbp2x*, and shares a similarly broad MIC distribution (although with typically lower MIC values) [37, 40, 41]. Our data also show that mutations in *pbp2x* alone have small effects (Figure S2), while genotypes carrying four or more variants have the highest mean MIC (Figure S2). Our explanatory power increases greatly when we predict beta-lactam MIC from the total number of candidate variants carried (Figure S10b). Although official breakpoints are lacking for many beta-lactam antibiotics in *S. suis*, for penicillin, only the isolates with the most variants reach clinical significance (i.e. penicillin resistance ≥ 1 or log(0) MIC).

In *S. pneumoniae* it has also been noted that mutations conferring resistance to beta-lactams occur in a set order, with amino acid changes in PBP2B and PBP2X often acting as the first step [36, 42, 43]. Figure 4 shows that our data also show this characteristic “nested” pattern (see also Figure S12; [44]), with PBP2X mutations largely occurring in backgrounds containing PBP2B mutations and *mraY* mutations occurring in backgrounds containing both PBP2B and PBP2X mutations.

**Figure 4.**
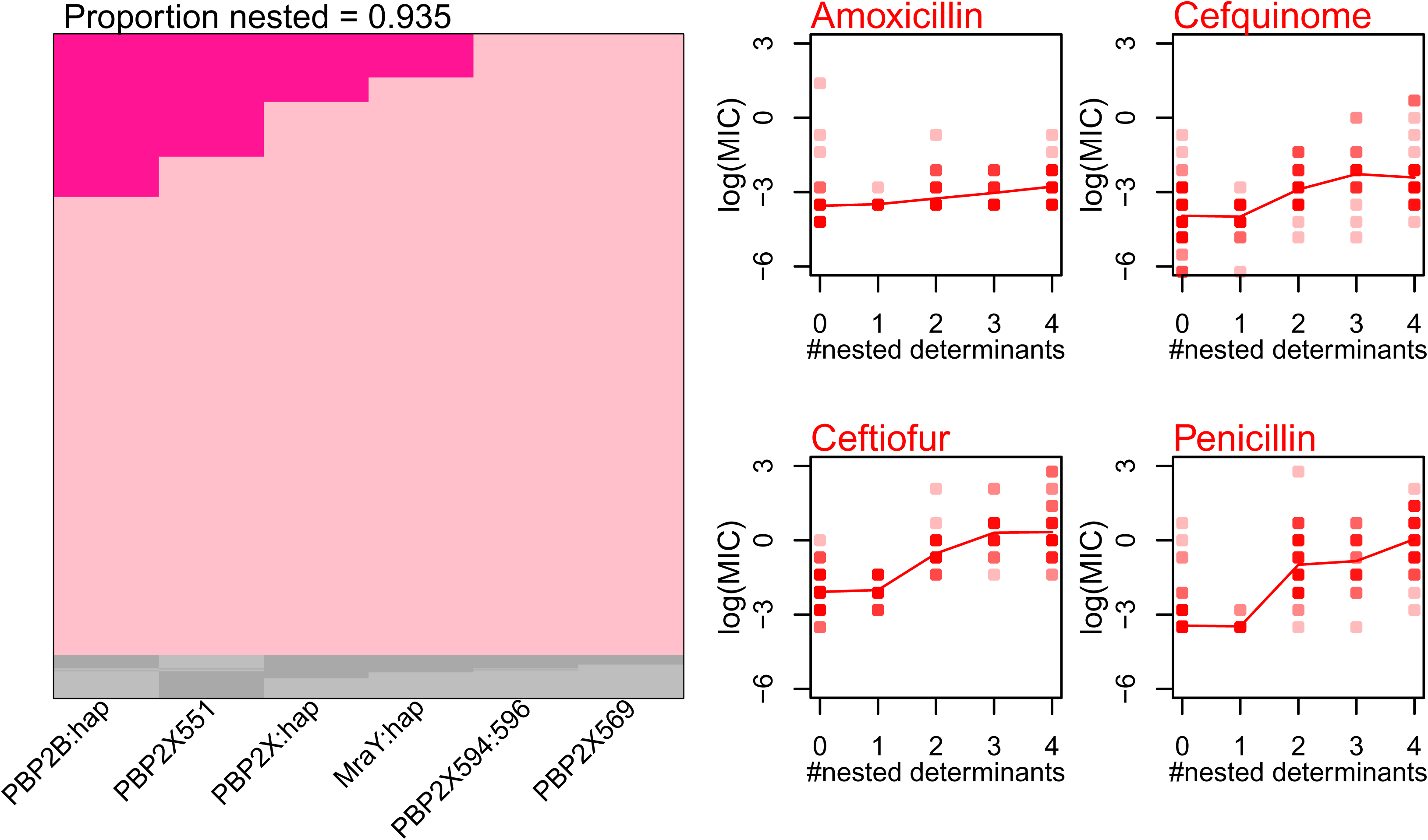
Beta-lactam resistance determinants have additive effects, and appear in a consistent order. The left-hand panel shows that the candidate determinants against the beta-lactam drug class often appear in a consistent order. For example, the mutation at site 551 of PBP2X tends to appear in backgrounds that already carry mutations in PBP2B. The plot was generated according to the method and plotting convention of [44], where each row represents an isolate, and those isolates that fit the nested pattern are shown in pink. Results show that 93.5% (634/678) of our isolates fit the nested pattern. The right-hand plots show the log MIC values for these 634 isolates, comparing isolates carrying different numbers of candidate determinants.

It is notable that mutations in *mraY* – which are generally last to occur, and never occur alone – might have a compensatory effect. Altered PBPs, while conferring resistance, might be less active transpeptidases than their wild type counterparts [45]. For example, the amidotransferase enzyme encoded by *murT* enables cross-linking of cell wall peptidoglycans by preparing lipid II used by pneumococcal PBPs [46, 47]. Mutations in *mraY* might therefore compensate for reduced enzymatic activity of altered PBPs.

Studies have shown that cefotaxime, a third-generation cephalosporin like ceftiofur, selectively inactivates PBP2X but not PBP2B [48] and that mutations within the catalytic domain in PBP2X give rise to cefotaxime, but not to penicillin, resistance [49]. Our results echo this pattern because our PBP2B haplotype has a smaller change on mean MIC of ceftiofur than penicillin (Figure S2), and the explanatory effects of our PBP haplotypes differ between the drugs (Figure 1). Previous variants in PBP2X thought to confer cefuroxime resistance [28], whilst validated by our data, were only found at very low frequency in our collection (Table 2).

## Discussion

We have examined antimicrobial susceptibility of *Streptococcus suis* to 16 antibiotics that are commonly used in pigs. Replicated measures of MIC suggest that, even when some isolates have very high MIC, there is often no natural separation into “resistant” and “susceptible” (Figure 1). This highlights both the need for clinical breakpoints in *S. suis*, and the need to evaluate AMR with quantitative measurements of MIC. This is particularly important for beta-lactam resistance, given its multi-allelic nature (Figures 4, S2), and the possibility of “MIC creep” that may eventually lead to treatment failures (e.g. vancomycin resistance in MRSA [50]).

We have also shown that AMR levels differ systematically between countries. For example, compared to the UK, Canada has consistently higher MICs, and frequencies of the same candidate AMR determinants (Figures 2, S1). Differences in antimicrobial usage are the most plausible explanation for this trend, because usage is much higher in Canada than the UK [24, 25]. However, our results are correlational and other factors could be explanatory e.g. different microbiome compositions in pigs from different areas.

Next, we have identified 43 candidate resistance determinants, and shown that they explain the majority of the observed variation in MIC for 11/16 antibiotics, and a substantial fraction of the variation for a further 2/16 (Figure 1).

While many of these variants were previously described, and most were found at low frequency (Table 2; Figure 2), we detected three novel candidates that are common and associated with clinically-relevant changes in MIC: *vgaF*, DHFR102 and the *dhfr* promoter. A new mechanism for tiamulin resistance, such as the *vgaF*, could have implications for treatment success because tiamulin is a common drug for treating infections in livestock, including pigs [51]. High-level trimethoprim resistance was associated with mutations in the promoter regions of the *dhfr* gene. This is in contrast to the *folA* (I100L), *folP* 1-2 codon insertion combination conferring high level resistance to trimethoprim/sulfamethoxazole (TMS) in *S. pneumoniae* [52]. While our MICs were for trimethoprim, our observations suggest the evolution of divergent mechanisms of resistance against TMS between these species. If these resistance mechanisms are functionally verified, they should be included in routine AMR gene testing for *S. suis*.

While our candidate variants were explanatory, we also found several complexities in the genotype-phenotype map. These include epistatic interactions, where the effects of some candidate genes vary with their *S. suis* genetic background. Phenotypic reversion (i.e. mutations in transcriptional regulators or elsewhere in the genome, counteracting a resistance determinant) can be common if resistance has a fitness cost [53], and this could explain why we see many isolates carrying candidate determinants but with low MIC (Figure 1). This same pattern is also apparent in up to 10% of *S. aureus* isolates [54]. These complexities have consequences for diagnostic investigation of AMR of *S. suis* using whole genome sequences. While the predictive power is quite high for many antibiotics, we should expect many Type I errors (false positives). These results also caution against using the presence of AMR determinants as a measure of resistance more generally.

While susceptibility to penicillin in clinical cases of *S. suis* remains high, we see substantial variation in beta-lactam MIC (Figure 1a). Consistent with studies in other Streptococci, our results show incremental and ordered changes in amino acids, first within PBP2A and then PBP2X that lead to clinically relevant elevations in MICs. Accurate predictions of genotype-phenotype have already been developed in *S. pneumoniae* [37, 52, 55–57], group B Streptococci [58] and *S. pyogenes* [31]. With no effective treatment for *S. suis* other than antibiotics, developing similar models and monitoring of these variants in *S. suis* populations should be a priority.

Two final aspects of our results highlight the fact that AMR is not a problem that can be tackled one disease at a time.

First, we have found high levels of resistance in *S. suis* even to antibiotics that are not typically used to treat this infection, including high rates of multi-drug resistance (Figure 3). This trend could be partly due to the co-occurrence of resistance genes on MGEs. However, determinants for beta lactams (found in 190/678 isolates) – which are the primary treatment – were less common than those for tetracyclines (574/678), MLS_B_ (395/678), trimethoprim (246/678) and tiamulin (193/678). This pattern probably reflects a “bystander selection effect”, common in opportunistic pathogens that are part of the healthy microbiota and frequently exposed to antibiotics used as growth promoters or therapeutics.

Second, many of the rarer AMR genes in our sample were not previously known in *S. suis*, but are common in other bacteria, both gram negative and gram positive (Table 2). Together with the presence of non-typical Streptococcal *ermB* in some isolates, this raises the possibility of *S. suis* acting as a reservoir for AMR determinants in other bacteria. Indeed *S. suis* shares conserved chromosomal insertion sites with many human pathogens, such as *S. pyogenes, S. pneumoniae*, and *S. agalactiae* [17]. Given that *S. suis* causes human clinical disease, a direct exchange of AMR genes between *S. suis* and other human pathogens is plausible. Our results lend further support to a one health approach to tackling AMR.

## Material and Methods

### Streptococcus suis isolates

We assembled a collection of 678 *S. suis* isolates, including isolates from the UK (*n*=423), Canada (*n*=205) and Vietnam (*n*=50) (Table S1).

In the UK, isolates came from three different collections. The first collection in 2009-2011 sampled non-clinical and clinical isolates from pigs across England and Wales (described in Weinert *et al.* 2015 [20]). The second collection in 2013-2014 sampled non-clinical isolates from five farms (described in Zou *et al.* 2018 [22]). The third collection during 2013–2014 targeted clinical isolates from pigs across England and Wales (described in Wileman *et al.* 2019 [21]). In pigs that showed clinical symptoms consistent with *S. suis* infections (e.g. meningitis, septicaemia and arthritis), the site of isolation was classified as ‘systemic’ if recovered from systemic sites. The site of recovery was classified as ‘respiratory’ if derived from lungs with gross lesions of pneumonia. *S. suis* isolates from the tonsils or tracheo-bronchus of healthy pigs or dead pigs without any typical signs of *S. suis* infections were defined as ‘non-clinical’. Isolates that could not confidently be assigned to these categories (e.g. a tonsil isolate from a pig with systemic signs) were classified as unknown. Altogether, the UK isolates were classified by clinical status as ‘systemic’ (*n*=94), ‘respiratory’ (*n*=50), ‘non-clinical’ (*n*=197) or ‘unknown’ (*n*=82), respectively.

The Canadian pig *S. suis* isolates from 1983-2016 were collected to target similar numbers of clinical and non-clinical isolates and were also classified by clinical status as ‘systemic’ (*n*=81), ‘respiratory’ (*n*=30), ‘non-clinical’ (*n*=55) or ‘unknown’ (*n*=39).

The Vietnamese isolates were collected to sample related populations from human and pig (described in Weinert *et al.* 2015 [20]). These comprised ‘systemic’ isolates (*n*=28) from human clinical cases of meningitis from provinces in southern and central Vietnam, and ‘systemic’ (*n*=4) or ‘non-clinical’ isolates (*n*=18) from pigs, collected between 2000 and 2010. These isolates were exclusively serotype 2 or serotype 14 and belong to one genetic population (Table S1).

### Antimicrobial susceptibility testing

The minimum inhibitory concentrations (MIC) for a range of antibiotics were determined by the micro-broth dilution method, which was performed and the results interpreted in accordance with CLSI Approved Standards, M100-S25 (2015), Vet01S 3^rd^ Edition (2015) and de Jong *et al.* (2014) [9, 59]. MIC measurements for some of our isolates were previously published [9]. For the remaining MIC measurements, the MICs were determined for sixteen different antimicrobial compounds, representing nine antimicrobial classes at LGC, Fordham, UK (formerly Quotient Bioresearch, Fordham, UK) for the UK and the Canadian isolates. MIC testing of antibiotics for the Vietnamese isolates was performed at the Oxford University Clinical Research Unit, Ho Chi Minh City, Vietnam, in collaboration with the Department of Veterinary Medicine, University of Cambridge, UK.

### Whole genome sequencing, assembly and inference of population structure

Genome DNA extractions and whole genome sequencing of newly sequenced *S suis* in this study were as previously described by Weinert *et al.* (2015) [20]. Briefly, single colonies of strains were grown up in broth culture, DNA was extracted using DNeasy kits (Qiagen), Illumina library preparations were performed as described by Quail *et al.* 2008 [60] and the whole genomes sequenced on the HiSeq2000 according to the manufacturer’s instructions (Illumina, San Diego, CA, USA) at the Welcome Trust Sanger Institute, Cambridge, UK. Sequencing generated 125bp paired end reads, which were assessed for quality using Sickle [61] after the removal of adapter sequences.

Sequencing reads that passed the quality threshold were put forward for *de-novo* assembly generation using Spades v.3.10.1 [62] utilising a variety of parameters conditions. Our 678 isolates were combined with 401 additional genomes given in Table S3 to increase robustness of *S. suis* population structure estimation, although these isolates were unavailable for MIC testing. Draft genomes were annotated using Prokka v1.12-beta (v2.8.2). [63] and bacterial species assignment was performed by a combination of MLST assignment and FastQ Screen v.0.11.1 using a custom database [64]. We mapped the Illumina reads back to the *de-novo* assembly to investigate polymorphic reads in the samples (indicative of mixed cultures) using BWA v.0.7.16a [65]. Genomes that exhibited poor sequencing quality (i.e poor assembly as indicated by a large number of contigs, low N50 values or a high number of polymorphic reads) or that which were inconsistent with an *S. suis* species assignment were excluded from the analysis.

In order to group our *S. suis* isolates in to different genetic clusters, we used the program hierBAPs in R [66]. First, we inferred core genes from our isolates using Roary (v2.8.2) [67], aligned them using MACSE [68], and stripped regions that could not be aligned unambiguously due to high divergence, indels or missing data. This conserved region of the core genome was used as input in hierBAPS.

### Known AMR determinant detection

ARIBA [69] identifies AMR determinants (or any sequence of interest) directly from paired sequencing reads using a public or custom reference database and relies on mapping the reads to reference sequence clusters followed by the formation of local assemblies of the mapped reads. The tool is further able to confirm the intactness of resistance genes to identify known or novel SNPs within a gene of interest.

The AMR determinants in *S. suis* were identified using ARIBA v2.10.0 based on the public AMR database, CARD [27], which was further supplemented with a custom database. The custom database contained gene sequences not present within the CARD database at the time of testing. These were previously published *S. suis* AMR genes, single nucleotide polymorphisms (SNPs) in genes known to confer antibiotic resistance and AMR genes found in other bacteria. In addition, we investigated whether there were additional novel resistance variants in known resistance genes (described below). The sequence identity threshold against a reference was set at 90%. Only paired-end sequence reads that passed the quality control thresholds were used as input for ARIBA. To identify genes falling under the 90% identity threshold, we also performed blast searches of the draft assemblies against the non-redundant NCBI protein or nucleotide databases to identify variants or chimeric alleles of known resistance determinants that might not be present in the CARD database. Examples of allelic variants identified this way include *tet*, *aad*, *ant* and *cat* (Table 2).

### Novel AMR candidate determinant detection

We identified novel resistance determinants by a range of methods. Candidate determinants that might encode novel resistance mechanisms were identified by scanning literature describing experimental studies and others found by Genome Wide Association Studies (GWAS) in related bacterial species, for example, variants of *pbp*s, *mraY*, *dhfr* and *folA* [36, 70–72] along with their promoter regions. To avoid over-fitting our generalised linear models, gene or promoters of interest in a “training” subset of the collection (*n*=205) were then extracted, aligned and ranked from the highest to lowest MIC values, using MUSCLE [73] in SEAVIEW [74]. Manual sequence analysis was then performed to identify either amino acid or nucleotide variations that associated with high MIC.

Kinetic and structural studies have previously established that beta-lactam resistance is conferred by substitutions within PBPs in *Streptococcus pneumoniae* [36, 75] and other streptococci [39, 76, 77]. While mutations are present throughout the entire PBPs, we noted statistically significant mutations (Table S2) in altered PBP2B and PBP2X proteins in strains exhibiting high penicillin (≥1mg/L) and ceftiofur MICs (≥2mg/L), respectively. In PBP2B, the residue variations were present in loop regions, K479T/A, D512E/Q/K/A, K513E/D and T515S, within the *S. suis* transpeptidase domain (residues 351-681). In PBP2X, the mutations were located at positions M437L, S445T, T467S, Y525F and T551S, also in loop regions within its catalytic domain (residues 265–619). No statistically significant variations were found within PBP1A, although the variant P405T was shared by some of the BAPs clusters. As multiple amino acid variants were identified within PBP2X and PBP2B, many of which were in the same genomic region (Table 2), we grouped the variants in to haplotypes. We did this by scoring haplotype presence if the isolate had all of the amino acid variants associated with high MIC in a given gene.

In addition to allelic differences in PBP2B and PBP2X, we also detected mutations in *mraY*. Found immediately upstream of *pbp2X*, *mraY* encodes an acetylmuramoyl-pentapeptide-transferase enzyme which is essential to lipid cycle reactions in the peptidoglycan cell wall biosynthesis pathway. Residue substitutions were present at A4S/T, M6I/L and G8S within the signal peptide regions of the protein. These substitutions were in strong linkage disequilibrium, such that they almost always occurred in pairs. In particular, both A4S/T and G8S occurred only in isolates also carrying M6I/L, while M6I/L appeared alone in only a single isolate. As such, we scored each isolate as carrying the *mraY* determinant only if it carried two of these three amino acid substitutions (either A4S/T and M6I/L, or G8S and M6I/L; Table 2). This set of isolates differed significantly in their MICs in both penicillin and ceftiofur across multiple BAPs groups (Table S2).

Not all isolates with high tiamulin MICs possessed a *lsaE* gene, suggesting an additional mechanism in *S. suis* conferring resistance to pleuromutilins. Using the CARD RGI (resistance gene identifier) protein homology models, a *vgaC* homologue was identified as a candidate gene associated with variation in tiamulin MIC. The gene, designated as *vgaF* in this work, encodes an ABC-F ATP-binding cassette ribosomal protection protein and shares 37.14% homology to the reference. Located within the chromosome, *vgaF* is found intact (∼1386bp) in statistically significant numbers in isolates with MIC≥ 8mg/L (Table S2) but is truncated (<750bp) in others. Plasmid borne *vga* homologues including *vgaC* are frequently detected in staphylococci and have been shown to confer cross resistance to pleuromutilins, lincosamides and streptogramin A antibiotics. A chromosome based *vgaA* gene variant, encoding an ATP-binding cassette protein conferring resistance to streptogramin A and related antibiotics in *S. aureus* has also been described [78].

An amino acid substitution (I102-L) was detected in the dihydrofolate reductase (DHFR) gene, *dhfr* (counterpart of *folA* in *S. pneumoniae*), in the majority of isolates with trimethoprim MICs of 0.12mg/L or greater. Similar substitutions of isoleucine to leucine at position 100 in *S. pneumoniae* [71, 79] and *Streptococcus pyogenes* [80] is known to cause resistance to TMP. Additionally, we also identified polymorphisms within the promoter region (0-30bp upstream) of the *dhfr* gene; an A5G substitution and insertions, in isolates exhibiting MICs≥1mg/L.

Mutations such as 1-2 codon insertions within *folP*, another core metabolic gene that is documented to confer resistance to trimethoprim/sulfamethoxazole in *S. pneumoniae*, were also examined. Whilst insertions and or deletions were absent, amino acid residue variations were observed at position 198 (A198G) either alone (9/678) or in combination with *dhfr* I102L (26/678 isolates) in isolates exhibiting a wide range of trimethoprim MICs; 0.03mg/L-32mg/mL. However, this data was not statistically significant in the Binomial sign tests, and hence was excluded from further analysis.

All of these variants might be causal, compensatory or linked to high MIC because of population structure. We therefore examined our variants and showed that susceptible and resistance alleles were found repeatedly in different BAPs clusters, which were significant using binomial sign tests when defining a cut-off (Table S2).

The *pbp*, *mraY* and *vgaF* genes were unable to be included into the custom database in ARIBA because of the sequence divergence. Therefore, we manually aligned the genes in all 678 isolates using MUSCLE [73] in SEAVIEW [74].

Similarly, analysis for known variants in *gyrA* and *parC* were also performed by manually aligning the genes. The *tet(O/W/32/O)* gene was scored by using blastn because ARIBA was unable to differentiate mosaic sequence patterns

### Serotype inference from whole genomes

Serotypes were determined *in silico* using the Athey *et al.* (2016) [81] serotype database, which we implemented in ARIBA. Failed ARIBA runs and sequence non-matches were designated as not available (NA).

### Statistical analyses

All general linear models were fit in R v. 3.3 [82] using the built-in function *lm* for models with solely fixed effects, or via Reduced Maximum Likelihood, using the function *lme* from the package nlme v. 3.1-141 [83], when BAPs cluster was included as a random effect. The response MIC values were log transformed. Our data for host and country is confounded because all of our human samples came from Vietnam (human *S. suis* has a higher prevalence in South East Asia [84]). Therefore, we coded “Country” as separate populations – Canada, UK, Vietnam-pig and Vietnam-human. We classified “serotypes” into either disease-associated or non-disease associated according to Wileman *et al.* (2019) [21].

### Data availability

Whole genome sequence assemblies are available from NCBI under Project ID PRJNA628943. The gene *cat* from isolate FX419 has accession MT367165. The gene *aadE1* from isolate CF2D3-1A has accession MT383663. The gene *aadE2* from isolate BH3D7-4E has accession MT383664. The gene *ant1* from isolate CF2D3-1C has accession MT383665. The gene *tetM* from isolate 1129711 has accession MT383666. The gene *tetM* from isolate LSS85 has accession MT383667. The gene *tetM* from isolate SS981 has accession MT383668. The gene tetO from isolate D16-010262 has accession MT383669. The gene *vgaF* from isolate 1230091 has accession MT431628.

## Acknowledgements

This work was primarily supported by a Challenge grant from the Royal Society (reference: CH160114) and an Isaac Newton Trust Research Grant (reference: 17.24(u)). In addition, LAW, ELM, GGRM were supported by a Sir Henry Dale Fellowship jointly funded by the Wellcome Trust and the Royal Society (109385/Z/15/Z). We also acknowledge a BBSRC grant to AWT (BBSRC grant BB/L003902/1), a LoLa grant to DJM and AWT (BB/G019274/1) and a China MoST grant to RZ (2013DFG32360) for sample collections. In addition, we wish to thank Abiyad Baig, Andrew Balmer and Olivier Restif for helpful discussion on antimicrobial resistance in *S. suis*.

## Contributions

LAW and NTH conceived the study. AWT, SMW, RZ, MG, NF, TMW, JP, DJM contributed isolates. NFH, ELM, PLKY, HDP, NTH collected new data. NFH, ELM, GGRM, JJW and LAW analysed data. JJW produced all figures. LAW, NFH and JJW wrote the manuscript.

## References

1. de Moor CE: Septicaemic infections in pigs, caused by haemolytic streptococci of new Lancefield groups designated R, S and T. Antonie van Leeuwenhoek 1963, 29(1):272–280.

2. Wertheim HFL, Nguyen HN, Taylor W, Lien TTM, Ngo HT, Nguyen TQ, Nguyen BNT, Nguyen HH, Nguyen HM, Nguyen CT et al: Streptococcus suis, an Important Cause of Adult Bacterial Meningitis in Northern Vietnam. PloS one 2009, 4(6):e5973.

3. Baums CG, Kock C, Beineke A, Bennecke K, Goethe R, Schröder C, Waldmann K-H, Valentin-Weigand P: Streptococcus suis bacterin and subunit vaccine immunogenicities and protective efficacies against serotypes 2 and 9. Clinical and vaccine immunology 2009, 16(2):200–208.

4. van Rennings L, von Münchhausen C, Ottilie H, Hartmann M, Merle R, Honscha W, Käsbohrer A, Kreienbrock L: Cross-sectional study on antibiotic usage in pigs in Germany. PloS one 2015, 10(3):e0119114.

5. McGlone JJ: The Future of Pork Production in the World: Towards Sustainable, Welfare-Positive Systems. Animals 2013, 3(2):401–415.

6. Steinfeld H, Gerber PJ, Wassenaar T, Castel V, Rosales M, De haan C: Livestock’s Long Shadow: Environmental Issues and Options, vol. 24; 2006.

7. Goyette-Desjardins G, Auger J-P, Xu J, Segura M, Gottschalk M: Streptococcus suis, an important pig pathogen and emerging zoonotic agent—an update on the worldwide distribution based on serotyping and sequence typing. Emerging Microbes and Infections 2014, 3(1):1–20.

8. Van Boeckel TP, Brower C, Gilbert M, Grenfell BT, Levin SA, Robinson TP, Teillant A, Laxminarayan R: Global trends in antimicrobial use in food animals. Proceedings of the National Academy of Sciences 2015, 112(18):5649–5654.

9. Hernandez-Garcia J, Wang J, Restif O, Holmes MA, Mather AE, Weinert LA, Wileman TM, Thomson JR, Langford PR, Wren BW et al: Patterns of antimicrobial resistance in Streptococcus suis isolates from pigs with or without streptococcal disease in England between 2009 and 2014. Vet Microbiol 2017, 207:117–124.

10. Yongkiettrakul S, Maneerat K, Arechanajan B, Malila Y, Srimanote P, Gottschalk M, Visessanguan W: Antimicrobial susceptibility of Streptococcus suis isolated from diseased pigs, asymptomatic pigs, and human patients in Thailand. BMC Veterinary Research 2019, 15(1):5.

11. Marie J, Morvan H, Berthelot-Hérault F, Sanders P, Kempf I, Gautier-Bouchardon AV, Jouy E, Kobisch M: Antimicrobial susceptibility of Streptococcus suis isolated from swine in France and from humans in different countries between 1996 and 2000. Journal of Antimicrobial Chemotherapy 2002, 50(2):201–209.

12. Šeol B, Kelnerić Ž, Hajsig D, Madić J, Naglić T: Susceptibility to Antimicrobial Agents of Streptococcus suis Capsular Type 2 Strains Isolated from Pigs. Zentralblatt für Bakteriologie 1996, 283(3):328–331.

13. Turgeon PL, Higgins R, Gottschalk M, Beaudoin M: Antimicrobial susceptibility of Streptococcus suis isolates. British Veterinary Journal 1994, 150(3):263–269.

14. Gurung M, Tamang MD, Moon DC, Kim S-R, Jeong J-H, Jang G-C, Jung S-C, Park Y-H, Lim S-K: Molecular Basis of Resistance to Selected Antimicrobial Agents in the Emerging Zoonotic Pathogen Streptococcus suis. Journal of Clinical Microbiology 2015, 53(7):2332–2336.

15. Yao J, Shang K, Huang J, Ran W, Kashif J, Wang L: Overexpression of an ABC transporter and mutations of GyrA, GyrB, and ParC in contributing to high-level ciprofloxacin resistance in Streptococcus suis type 2. BioScience Trends 2014, 8(2):84–92.

16. Hoa NT, Chieu TT, Nghia HD, Mai NT, Anh PH, Wolbers M, Baker S, Campbell JI, Chau NV, Hien TT et al: The antimicrobial resistance patterns and associated determinants in Streptococcus suis isolated from humans in southern Vietnam, 1997-2008. BMC infectious diseases 2011, 11:6.

17. Huang J, Ma J, Shang K, Hu X, Liang Y, Li D, Wu Z, Dai L, Chen L, Wang L: Evolution and Diversity of the Antimicrobial Resistance Associated Mobilome in Streptococcus suis: A Probable Mobile Genetic Elements Reservoir for Other Streptococci. Frontiers in Cellular and Infection Microbiology 2016, 6(118).

18. Hu P, Yang M, Zhang A, Wu J, Chen B, Hua Y, Yu J, Chen H, Xiao J, Jin M: Comparative genomics study of multi-drug-resistance mechanisms in the antibiotic-resistant Streptococcus suis R61 strain. PloS one 2011, 6(9):e24988.

19. Holden MT, Hauser H, Sanders M, Ngo TH, Cherevach I, Cronin A, Goodhead I, Mungall K, Quail MA, Price C et al: Rapid evolution of virulence and drug resistance in the emerging zoonotic pathogen Streptococcus suis. PloS one 2009, 4(7):e6072.

20. Weinert LA, Chaudhuri RR, Wang J, Peters SE, Corander J, Jombart T, Baig A, Howell KJ, Vehkala M, Välimäki N et al: Genomic signatures of human and animal disease in the zoonotic pathogen Streptococcus suis. Nature Communications 2015, 6(1):6740.

21. Wileman TM, Weinert LA, Howell KJ, Wang J, Peters SE, Williamson SM, Wells JM, Langford PR, Rycroft AN, Wren BW et al: Pathotyping the Zoonotic Pathogen Streptococcus suis: Novel Genetic Markers To Differentiate Invasive Disease-Associated Isolates from Non-Disease-Associated Isolates from England and Wales. Journal of Clinical Microbiology 2019, 57(7):e01712–01718.

22. Zou G, Zhou J, Xiao R, Zhang L, Cheng Y, Jin H, Li L, Zhang L, Wu B, Qian P et al: Effects of Environmental and Management-Associated Factors on Prevalence and Diversity of Streptococcus suis in Clinically Healthy Pig Herds in China and the United Kingdom. Applied and Environmental Microbiology 2018, 84(8):e02590–02517.

23. van Hout J, Heuvelink A, Gonggrijp M: Monitoring of antimicrobial susceptibility of Streptococcus suis in the Netherlands, 2013–2015. Vet Microbiol 2016, 194:5–10.

24. Dunlop RH, McEwen SA, Meek AH, Black WD, Clarke RC, Friendship RM: Individual and group antimicrobial usage rates on 34 farrow-to-finish swine farms in Ontario, Canada. Preventive Veterinary Medicine 1998, 34(4):247–264.

25. Directorate VM: UK veterinary antibiotic resistance and sales surveillance report. In.; 2018.

26. Carrique-Mas JJ, Choisy M, Van Cuong N, Thwaites G, Baker S: An estimation of total antimicrobial usage in humans and animals in Vietnam. Antimicrobial Resistance and Infection Control 2020, 9(1):16.

27. Jia B, Raphenya AR, Alcock B, Waglechner N, Guo P, Tsang KK, Lago BA, Dave BM, Pereira S, Sharma AN, et al: CARD 2017: expansion and model-centric curation of the comprehensive antibiotic resistance database. Nucleic Acids Res 2017, 45(D1):D566–D573.

28. Ge Y, Wu J, Xia Y, Yang M, Xiao J, Yu J: Molecular dynamics simulation of the complex PBP-2x with drug cefuroxime to explore the drug resistance mechanism of Streptococcus suis R61. PloS one 2012, 7(4):e35941.

29. Palmieri C, Magi G, Mingoia M, Bagnarelli P, Ripa S, Varaldo PE, Facinelli B: Characterization of a Streptococcus suis tet(O/W/32/O)-carrying element transferable to major streptococcal pathogens. Antimicrobial agents and chemotherapy 2012, 56(9):4697–4702.

30. Escudero JA, San Millan A, Catalan A, de la Campa AG, Rivero E, Lopez G, Dominguez L, Moreno MA, Gonzalez-Zorn B: First characterization of fluoroquinolone resistance in Streptococcus suis. Antimicrobial agents and chemotherapy 2007, 51(2):777–782.

31. Vannice KS, Ricaldi J, Nanduri S, Fang FC, Lynch JB, Bryson-Cahn C, Wright T, Duchin J, Kay M, Chochua S et al: Streptococcus pyogenes pbp2x Mutation Confers Reduced Susceptibility to β-Lactam Antibiotics. Clinical infectious diseases : an official publication of the Infectious Diseases Society of America 2020, 71(1):201–204.

32. Chewapreecha C, Marttinen P, Croucher NJ, Salter SJ, Harris SR, Mather AE, Hanage WP, Goldblatt D, Nosten FH, Turner C et al: Comprehensive identification of single nucleotide polymorphisms associated with beta-lactam resistance within pneumococcal mosaic genes. PLoS genetics 2014, 10(8):e1004547.

33. Albarracin Orio AG, Pinas GE, Cortes PR, Cian MB, Echenique J: Compensatory evolution of pbp mutations restores the fitness cost imposed by beta-lactam resistance in Streptococcus pneumoniae. PLoS pathogens 2011, 7(2):e1002000.

34. Leclercq R: Mechanisms of Resistance to Macrolides and Lincosamides: Nature of the Resistance Elements and Their Clinical Implications. Clinical Infectious Diseases 2002, 34(4):482–492.

35. Farrow KA, Lyras D, Polekhina G, Koutsis K, Parker MW, Rood JI: Identification of Essential Residues in the Erm(B) rRNA Methyltransferase of Clostridium perfringens. Antimicrobial agents and chemotherapy 2002, 46(5):1253.

36. Grebe T, Hakenbeck R: Penicillin-binding proteins 2b and 2x of Streptococcus pneumoniae are primary resistance determinants for different classes of beta-lactam antibiotics. Antimicrobial agents and chemotherapy 1996, 40(4):829–834.

37. Li Y, Metcalf BJ, Chochua S, Li Z, Gertz RE, Walker H, Hawkins PA, Tran T, Whitney CG, McGee L et al: Penicillin-Binding Protein Transpeptidase Signatures for Tracking and Predicting β-Lactam Resistance Levels in Streptococcus pneumoniae. mBio 2016, 7(3):e00756–00716.

38. Dahesh S, Hensler ME, Van Sorge NM, Gertz RE, Jr., Schrag S, Nizet V, Beall BW: Point mutation in the group B streptococcal pbp2x gene conferring decreased susceptibility to beta-lactam antibiotics. Antimicrobial agents and chemotherapy 2008, 52(8):2915–2918.

39. Fuursted K, Stegger M, Hoffmann S, Lambertsen L, Andersen PS, Deleuran M, Thomsen MK: Description and characterization of a penicillin-resistant Streptococcus dysgalactiae subsp. equisimilis clone isolated from blood in three epidemiologically linked patients. Journal of Antimicrobial Chemotherapy 2016, 71(12):3376–3380.

40. Holmer I, Salomonsen CM, Jorsal SE, Astrup LB, Jensen VF, Høg BB, Pedersen K: Antibiotic resistance in porcine pathogenic bacteria and relation to antibiotic usage. BMC Veterinary Research 2019, 15(1):449.

41. Liñares J, Ardanuy C, Pallares R, Fenoll A: Changes in antimicrobial resistance, serotypes and genotypes in Streptococcus pneumoniae over a 30-year period. Clinical Microbiology and Infection 2010, 16(5):402–410.

42. Mouz N, Di Guilmi AM, Gordon E, Hakenbeck R, Dideberg O, Vernet T: Mutations in the active site of penicillin-binding protein PBP2x from Streptococcus pneumoniae. Role in the specificity for beta-lactam antibiotics. The Journal of biological chemistry 1999, 274(27):19175–19180.

43. Nagai K, Davies TA, Jacobs MR, Appelbaum PC: Effects of amino acid alterations in penicillin-binding proteins (PBPs) 1a, 2b, and 2x on PBP affinities of penicillin, ampicillin, amoxicillin, cefditoren, cefuroxime, cefprozil, and cefaclor in 18 clinical isolates of penicillin-susceptible, -intermediate, and -resistant pneumococci. Antimicrobial agents and chemotherapy 2002, 46(5):1273–1280.

44. Lehtinen S, Blanquart F, Lipsitch M, Fraser C, with the Maela Pneumococcal Collaboration: On the evolutionary ecology of multidrug resistance in bacteria. PLoS pathogens 2019, 15(5):e1007763.

45. Calvez P, Breukink E, Roper DI, Dib M, Contreras-Martel C, Zapun A: Substitutions in PBP2b from beta-Lactam-resistant Streptococcus pneumoniae Have Different Effects on Enzymatic Activity and Drug Reactivity. Journal of Biological Chemistry 2017, 292(7):2854–2865.

46. Figueiredo TA, Sobral RG, Ludovice AM, Almeida JM, Bui NK, Vollmer W, de Lencastre H, Tomasz A: Identification of genetic determinants and enzymes involved with the amidation of glutamic acid residues in the peptidoglycan of Staphylococcus aureus. PLoS pathogens 2012, 8(1):e1002508.

47. Zapun A, Philippe J, Abrahams KA, Signor L, Roper DI, Breukink E, Vernet T: In vitro reconstitution of peptidoglycan assembly from the Gram-positive pathogen Streptococcus pneumoniae. ACS chemical biology 2013, 8(12):2688–2696.

48. Kocaoglu O, Tsui HC, Winkler ME, Carlson EE: Profiling of beta-lactam selectivity for penicillin-binding proteins in Streptococcus pneumoniae D39. Antimicrobial agents and chemotherapy 2015, 59(6):3548–3555.

49. Coffey TJ, Daniels M, McDougal LK, Dowson CG, Tenover FC, Spratt BG: Genetic analysis of clinical isolates of Streptococcus pneumoniae with high-level resistance to expanded-spectrum cephalosporins. Antimicrobial agents and chemotherapy 1995, 39(6):1306–1313.

50. Sakoulas G, Moellering RC, Jr.: Increasing Antibiotic Resistance among Methicillin-Resistant Staphylococcus aureus Strains. Clinical Infectious Diseases 2008, 46(Supplement_5):S360–S367.

51. van Duijkeren E, Greko C, Pringle M, Baptiste KE, Catry B, Jukes H, Moreno MA, Pomba MCMF, Pyörälä S, Rantala M et al: Pleuromutilins: use in food-producing animals in the European Union, development of resistance and impact on human and animal health. Journal of Antimicrobial Chemotherapy 2014, 69(8):2022–2031.

52. Metcalf BJ, Chochua S, Gertz RE, Jr., Li Z, Walker H, Tran T, Hawkins PA, Glennen A, Lynfield R, Li Y et al: Using whole genome sequencing to identify resistance determinants and predict antimicrobial resistance phenotypes for year 2015 invasive pneumococcal disease isolates recovered in the United States. Clinical microbiology and infection : the official publication of the European Society of Clinical Microbiology and Infectious Diseases 2016, 22(12):1002.e1001-1002.e1008.

53. Dunai A, Spohn R, Farkas Z, Lázár V, Györkei Á, Apjok G, Boross G, Szappanos B, Grézal G, Faragó A et al: Rapid decline of bacterial drug-resistance in an antibiotic-free environment through phenotypic reversion. Elife 2019, 8:e47088.

54. Kime L, Randall CP, Banda FI, Coll F, Wright J, Richardson J, Empel J, Parkhill J, O’Neill AJ: Transient Silencing of Antibiotic Resistance by Mutation Represents a Significant Potential Source of Unanticipated Therapeutic Failure. mBio 2019, 10(5):e01755–01719.

55. Li Y, Metcalf BJ, Chochua S, Li Z, Gertz RE, Jr., Walker H, Hawkins PA, Tran T, McGee L, Beall BW: Validation of β-lactam minimum inhibitory concentration predictions for pneumococcal isolates with newly encountered penicillin binding protein (PBP) sequences. BMC genomics 2017, 18(1):621.

56. Beall B, Chochua S, Gertz RE, Jr., Li Y, Li Z, McGee L, Metcalf BJ, Ricaldi J, Tran T, Walker H et al: A Population-Based Descriptive Atlas of Invasive Pneumococcal Strains Recovered Within the U.S. During 2015-2016. Frontiers in microbiology 2018, 9:2670.

57. Varghese J, Chochua S, Tran T, Walker H, Li Z, Snippes Vagnone PM, Lynfield R, McGee L, Li Y, Metcalf BJ et al: Multistate population and whole genome sequence-based strain surveillance of invasive pneumococci recovered in the USA during 2017. Clinical microbiology and infection : the official publication of the European Society of Clinical Microbiology and Infectious Diseases 2020, 26(4):512.e511-512.e510.

58. Metcalf BJ, Chochua S, Gertz RE, Jr., Hawkins PA, Ricaldi J, Li Z, Walker H, Tran T, Rivers J, Mathis S et al: Short-read whole genome sequencing for determination of antimicrobial resistance mechanisms and capsular serotypes of current invasive Streptococcus agalactiae recovered in the USA. Clinical microbiology and infection : the official publication of the European Society of Clinical Microbiology and Infectious Diseases 2017, 23(8):574.e577-574.e514.

59. de Jong A, Thomas V, Simjee S, Moyaert H, El Garch F, Maher K, Morrissey I, Butty P, Klein U, Marion H et al: Antimicrobial susceptibility monitoring of respiratory tract pathogens isolated from diseased cattle and pigs across Europe: The VetPath study. Vet Microbiol 2014, 172(1):202–215.

60. Quail MA, Kozarewa I, Smith F, Scally A, Stephens PJ, Durbin R, Swerdlow H, Turner DJ: A large genome center’s improvements to the Illumina sequencing system. Nature Methods 2008, 5(12):1005–1010.

61. Joshi N, Fass J: Sickle: A sliding-window, adaptive, quality-based trimming tool for FastQ files. In.; 2011.

62. Bankevich A, Nurk S, Antipov D, Gurevich AA, Dvorkin M, Kulikov AS, Lesin VM, Nikolenko SI, Pham S, Prjibelski AD et al: SPAdes: a new genome assembly algorithm and its applications to single-cell sequencing. Journal of Computational Biology 2012, 19(5):455–477.

63. Seemann T: Prokka: rapid prokaryotic genome annotation. Bioinformatics 2014, 30(14):2068–2069.

64. Wingett SW, Andrews S: FastQ Screen: A tool for multi-genome mapping and quality control. F1000Res 2018, 7:1338–1338.

65. Li H, Durbin R: Fast and accurate short read alignment with Burrows– Wheeler transform. Bioinformatics 2009, 25(14):1754–1760.

66. Cheng L, Connor TR, Sirén J, Aanensen DM, Corander J: Hierarchical and Spatially Explicit Clustering of DNA Sequences with BAPS Software. Molecular biology and evolution 2013, 30(5):1224–1228.

67. Page AJ, Cummins CA, Hunt M, Wong VK, Reuter S, Holden MTG, Fookes M, Falush D, Keane JA, Parkhill J: Roary: rapid large-scale prokaryote pan genome analysis. Bioinformatics 2015, 31(22):3691–3693.

68. Ranwez V, Harispe S, Delsuc F, Douzery EJP: MACSE: Multiple Alignment of Coding SEquences Accounting for Frameshifts and Stop Codons. PloS one 2011, 6(9):e22594.

69. Hunt M, Mather AE, Sánchez-Busó L, Page AJ, Parkhill J, Keane JA, Harris SR: ARIBA: rapid antimicrobial resistance genotyping directly from sequencing reads. Microb Genom 2017, 3(10):e000131.

70. Lopez P, Espinosa M, Greenberg B, Lacks St: Sulfonamide resistance in Streptococcus pneumoniae: DNA sequence of the gene encoding dihydropteroate synthase and characterization of the enzyme. Journal of Bacteriology 1987, 169(9):4320–4326.

71. Adrian PV, Klugman KP: Mutations in the dihydrofolate reductase gene of trimethoprim-resistant isolates of Streptococcus pneumoniae. Antimicrobial agents and chemotherapy 1997, 41(11):2406–2413.

72. Hakenbeck R, Grebe T, Zähner D, Stock JB: β-Lactam resistance in Streptococcus pneumoniae: penicillin-binding proteins and non-penicillin-binding proteins. Molecular Microbiology 1999, 33(4):673–678.

73. Edgar RC: MUSCLE: multiple sequence alignment with high accuracy and high throughput. Nucleic Acids Res 2004, 32(5):1792–1797.

74. Gouy M, Guindon S, Gascuel O: SeaView version 4: A multiplatform graphical user interface for sequence alignment and phylogenetic tree building. Molecular biology and evolution 2010, 27(2):221–224.

75. Contreras-Martel C, Dahout-Gonzalez C, Martins Ados S, Kotnik M, Dessen A: PBP active site flexibility as the key mechanism for beta-lactam resistance in pneumococci. Journal of molecular biology 2009, 387(4):899–909.

76. Haenni M, Galofaro L, Ythier M, Giddey M, Majcherczyk P, Moreillon P, Madec JY: Penicillin-binding protein gene alterations in Streptococcus uberis isolates presenting decreased susceptibility to penicillin. Antimicrobial agents and chemotherapy 2010, 54(3):1140–1145.

77. Piccinelli G, Carlentini G, Gargiulo F, Caruso A, De Francesco MA: Analysis of Point Mutations in the pbp2x, pbp2b, and pbp1a Genes of Streptococcus agalactiae and Their Relation with a Reduced Susceptibility to Cephalosporins. Microbial Drug Resistance 2017, 23(8):1019–1024.

78. Haroche J, Allignet J, Buchrieser C, El Solh N: Characterization of a Variant of vga (A) Conferring Resistance to Streptogramin A and Related Compounds. Antimicrobial agents and chemotherapy 2000, 44(9):2271.

79. Pikis A, Donkersloot JA, Rodriguez WJ, Keith JM: A conservative amino acid mutation in the chromosome-encoded dihydrofolate reductase confers trimethoprim resistance in Streptococcus pneumoniae. Journal of Infectious Diseases 1998, 178(3):700–706.

80. Bergmann R, van der Linden M, Chhatwal GS, Nitsche-Schmitz DP: Factors that cause trimethoprim resistance in Streptococcus pyogenes. Antimicrobial agents and chemotherapy 2014, 58(4):2281–2288.

81. Athey TBT, Teatero S, Lacouture S, Takamatsu D, Gottschalk M, Fittipaldi N: Determining Streptococcus suis serotype from short-read whole-genome sequencing data. BMC Microbiol 2016, 16(1):162–162.

82. Team RC: R: A language and environment for statistical computing. 2013.

83. Pinheiro J, Bates D, DebRoy S, Sarkar D, Heisterkamp S, Van Willigen B, Maintainer R: Package ‘nlme’. 2019.

84. Palmieri C, Varaldo PE, Facinelli B: Streptococcus suis, an Emerging Drug-Resistant Animal and Human Pathogen. Frontiers in microbiology 2011, 2:235.

85. Huang K, Zhang Q, Song Y, Zhang Z, Zhang A, Xiao J, Jin M: Characterization of Spectinomycin Resistance in Streptococcus suis Leads to Two Novel Insights into Drug Resistance Formation and Dissemination Mechanism. Antimicrobial agents and chemotherapy 2016, 60(10):6390–6392.

86. Chang B, Wada A, Ikebe T, Ohnishi M, Mita K, Endo M, Matsuo H, Asatuma Y, Kuramoto S, Sekiguchi H et al: Characteristics of Streptococcus suis isolated from patients in Japan. Japanese journal of infectious diseases 2006, 59(6):397–399.

87. Bojarska A, Molska E, Janas K, Skoczynska A, Stefaniuk E, Hryniewicz W, Sadowy E: Streptococcus suis in invasive human infections in Poland: clonality and determinants of virulence and antimicrobial resistance. European Journal of Clinical Microbiology and Infectious Diseases 2016, 35(6):917–925.

88. Athey TB, Auger JP, Teatero S, Dumesnil A, Takamatsu D, Wasserscheid J, Dewar K, Gottschalk M, Fittipaldi N: Complex Population Structure and Virulence Differences among Serotype 2 Streptococcus suis Strains Belonging to Sequence Type 28. PloS one 2015, 10(9):e0137760.

