## Supplementary_information for "A comprehensive portrait of antimicrobial resistance in the zoonotic pathogen *Streptococcus suis*"

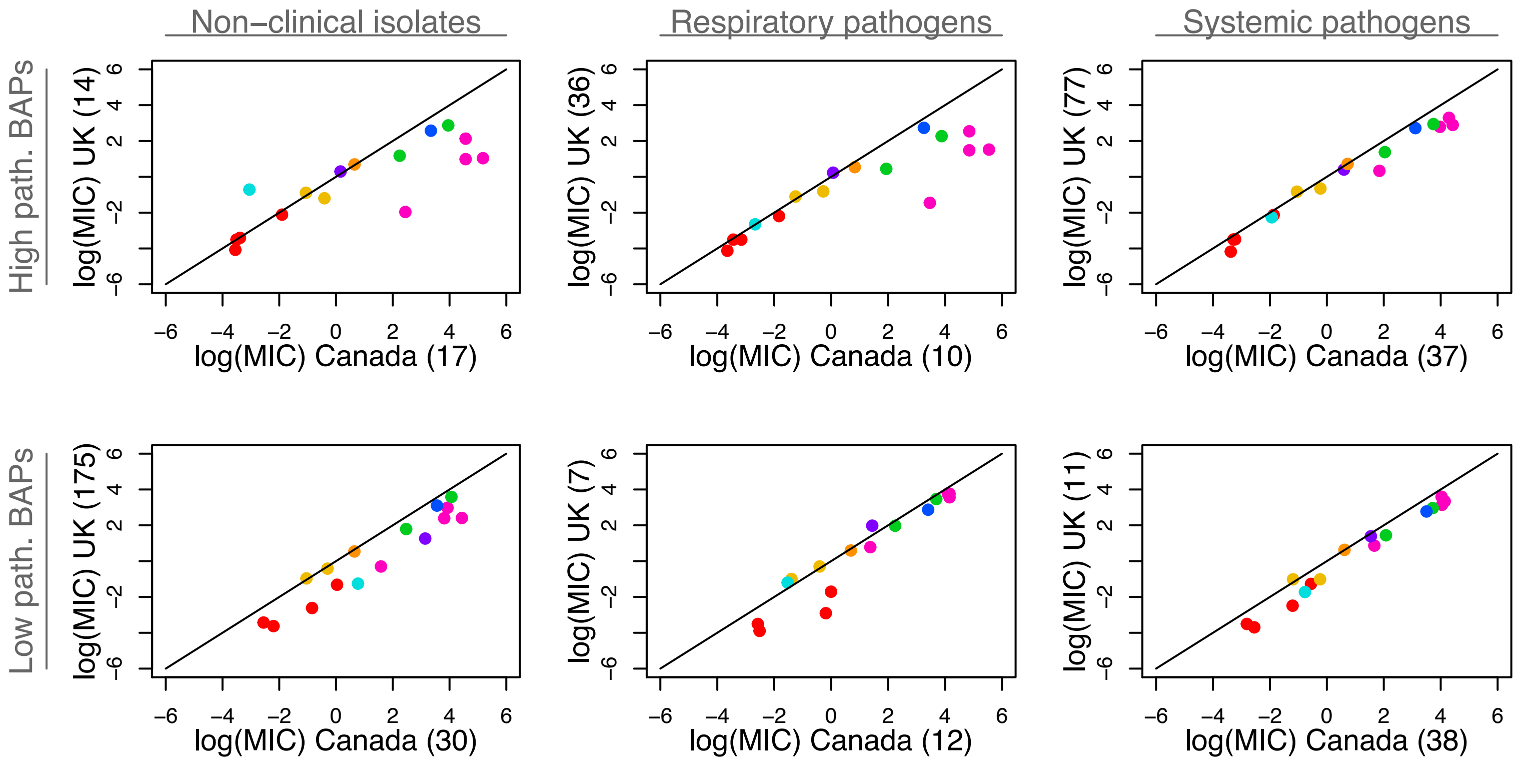

Non−clinical isolates
Respiratory pathogens
Systemic pathogens
High path. BAPs
Low path. BAPs
log(MIC) UK (14)
log(MIC) UK (36)
log(MIC) UK (77)
log(MIC) UK (175)
log(MIC) UK (7)
log(MIC) UK (11)
log(MIC) Canada (17)
log(MIC) Canada (10)
log(MIC) Canada (37)
log(MIC) Canada (30)
log(MIC) Canada (12)
log(MIC) Canada (38)
−6 −4 −2 0 2 4 6

**Figure S1. MIC values are consistently higher in Canada than the UK.** Each panel replicates the left-hand panel in Figure 2a but for a smaller subset of the isolates. In particular, the three columns contain results for non-clinical isolates from healthy pigs (left-hand column), respiratory pathogens (middle column) and systemic pathogens (right-hand columns). The two rows contain results for two groups of genetic clusters of *S. suis* as identified by [1]. The two groups are reciprocally monophyletic in a consensus phylogeny, and contain either a high proportion of disease-causing isolates (“High path. BAPs”) or a low proportion of disease-causing isolates (“Low path. BAPs”). The pattern of higher MICs in Canada is found consistently in all six subsets of the data.

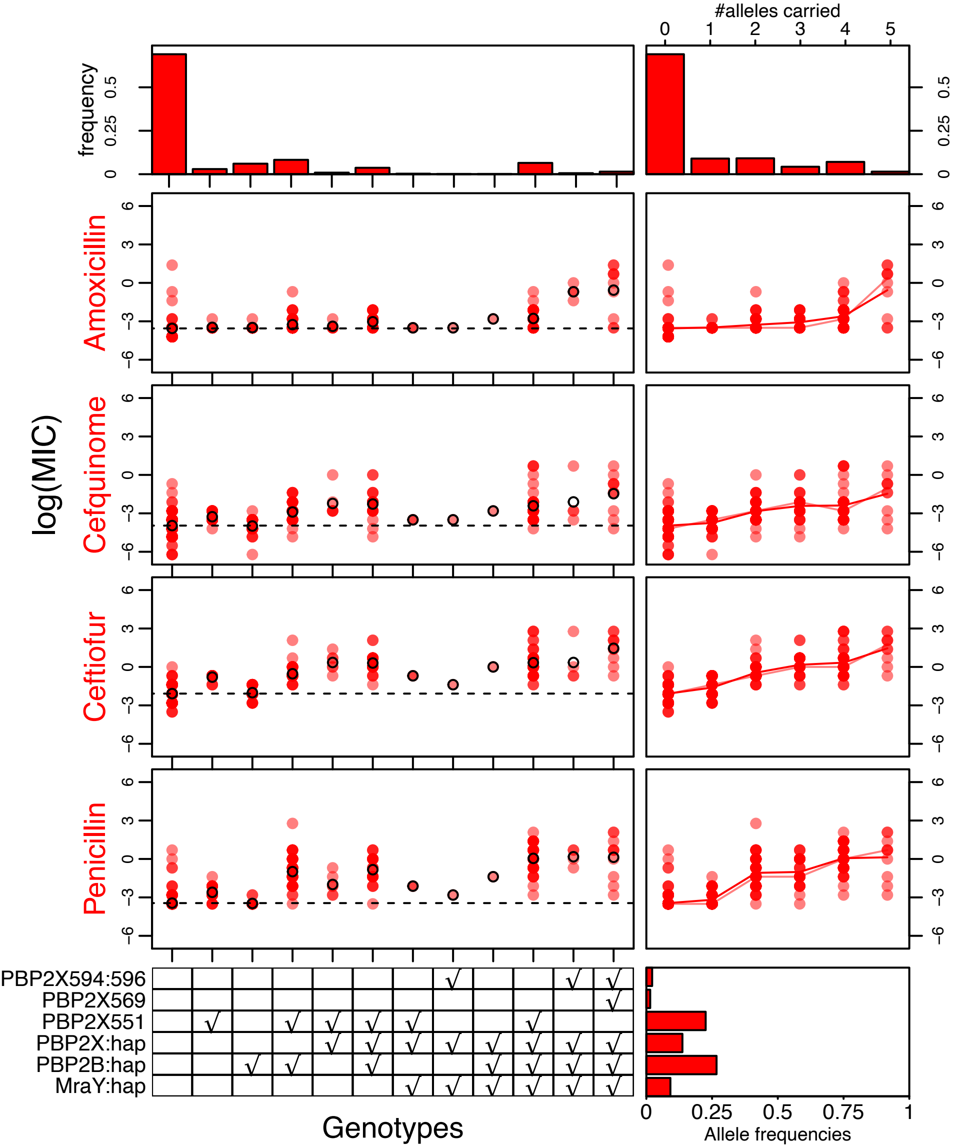
#alleles carried
frequency
log(MIC)
Amoxicillin
Cefquinome
Ceftiofur
Penicillin
PBP2X594:596
PBP2X569
PBP2X551
PBP2X:hap
PBP2B:hap
MraY:hap
Genotypes
Allele frequencies

**Figure S2. The effects of candidate determinants on MIC for beta-lactam antibiotics.** The left-hand panels show the log MIC values for each isolate containing a given combination of candidate resistance determinants to the beta-lactam class of antibiotics. In each case, the mean log MIC is shown by an empty circle, and the overall mean by a dashed line. The genotypes are indicated in the bottom panel, and the proportion of isolates carrying each genotype in the upper panel. The right-hand panels show results for isolates carrying different numbers of candidate resistance determinants, and the individual determinant frequencies. The cumulative effects of the determinants against beta-lactams are clearly visible from the right-hand plots.

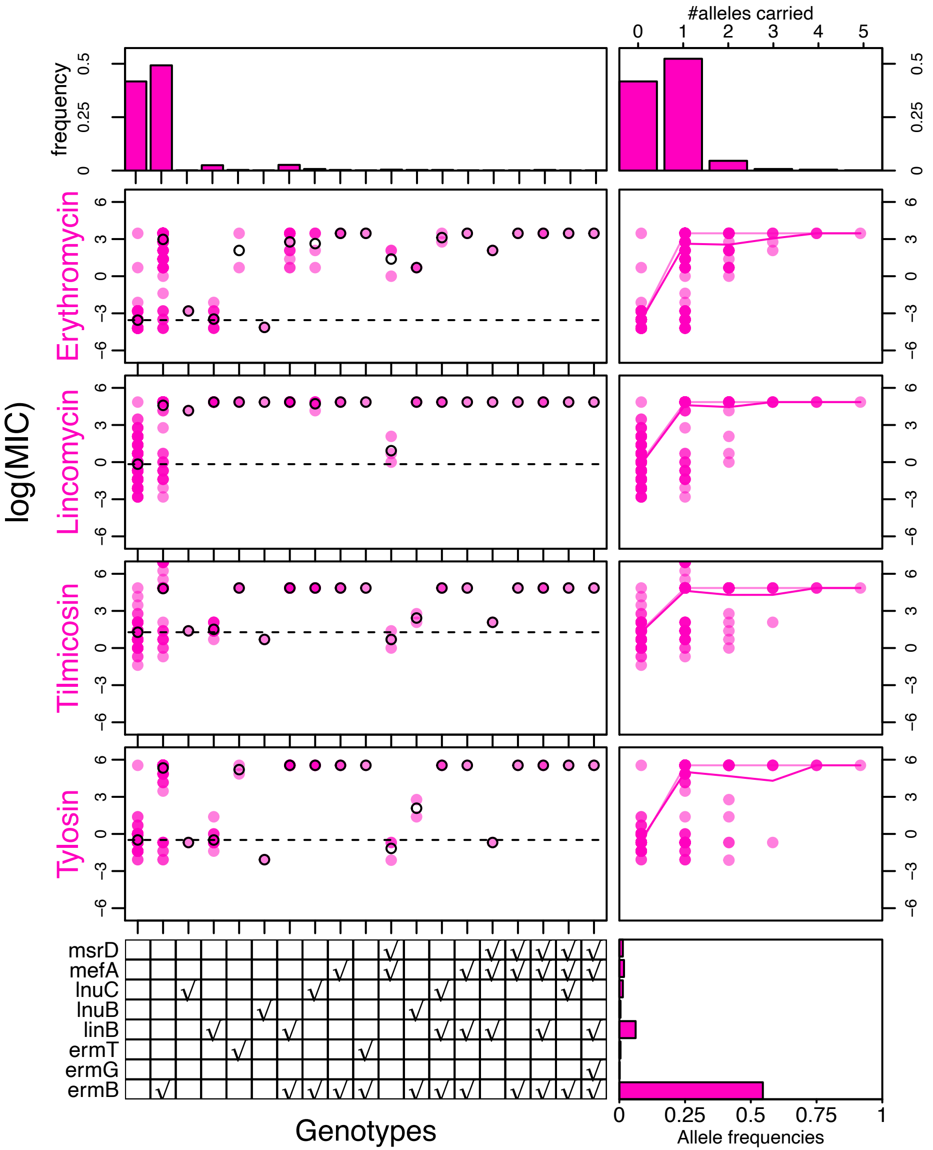

#alleles carried
0 1 2 3 4 5
frequency
0 0.25 0.5
log(MIC)
Erythromycin
Lincomycin
Tilmicosin
Tylosin
6 3 0 −3 −6
msrD
mefA
lnuC
lnuB
linB
ermT
ermG
ermB
Genotypes
0 0.25 0.5 0.75 1
Allele frequencies

**Figure S3. The effects of candidate determinants on MIC for macrolide-lincosamide-streptogramin B ($MLS_B$) antibiotics.** All details match Figure S2. As discussed in the text, determinants show an “all-or-nothing” effect, quite different to the pattern seen for the beta-lactams (Figure S2), while determinants such as *msrD* act only against the macrolide erythromycin, while others, such as *linB* act only against the three lincosamides.

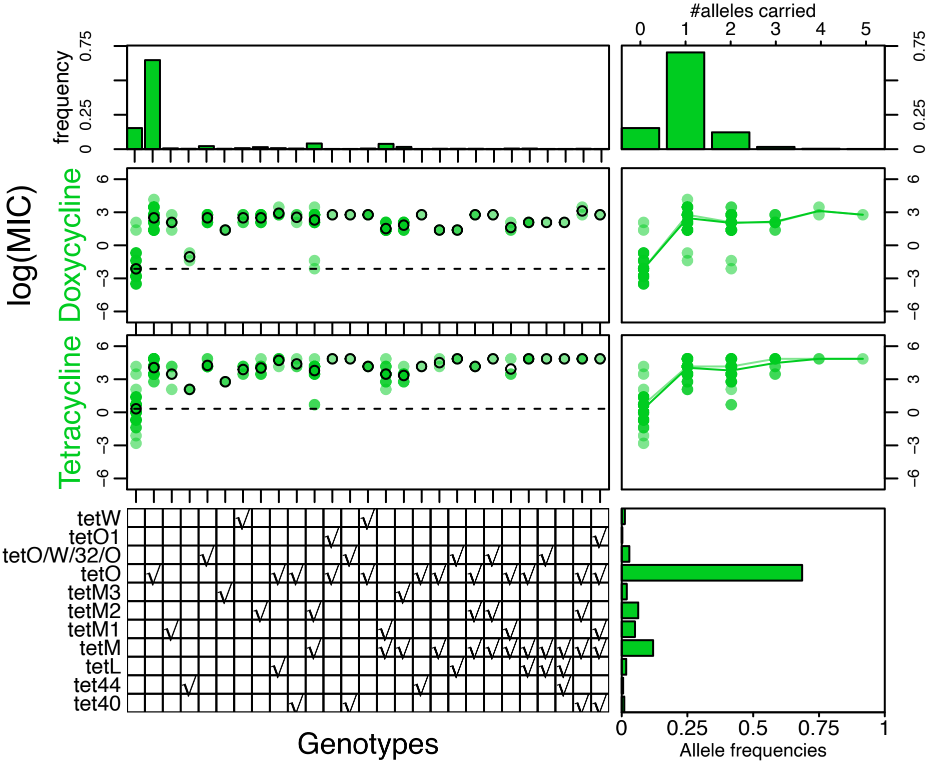
#alleles carried
0
1
2
3
4
5
frequency
0
0.25
0.75
log(MIC)
Doxycycline
Tetracycline
6
3
0
−3
−6
tetW
tetO1
tetO/W/32/O
tetO
tetM3
tetM2
tetM1
tetM
tetL
tet44
tet40
Genotypes
0
0.25
0.5
0.75
1
Allele frequencies

**Figure S4. The effects of candidate alleles on MIC for tetracycline antibiotics.** All details match Figure S2.

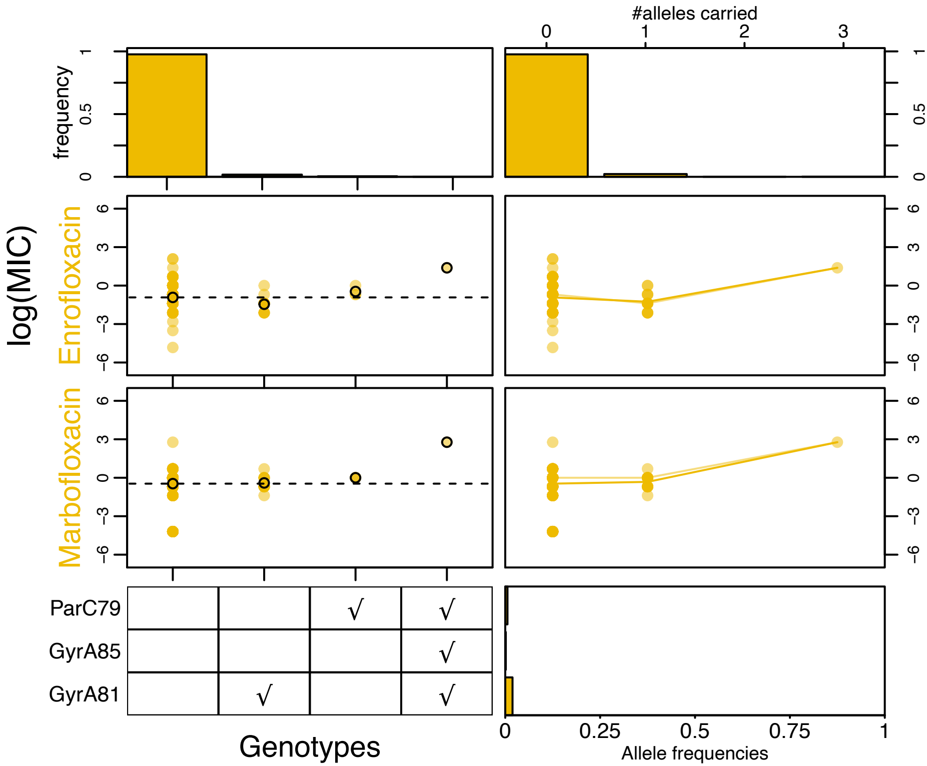

#alleles carried
0
1
2
3
frequency
0
0.5
1
log(MIC)
Enrofloxacin
Marbofloxacin
6
3
0
−3
−6
ParC79
GyrA85
GyrA81
√
Genotypes
0
0.25
0.5
0.75
1
Allele frequencies

**Figure S5. The effects of candidate alleles on MIC for fluoroquinolone antibiotics.** All details match Figure S2.

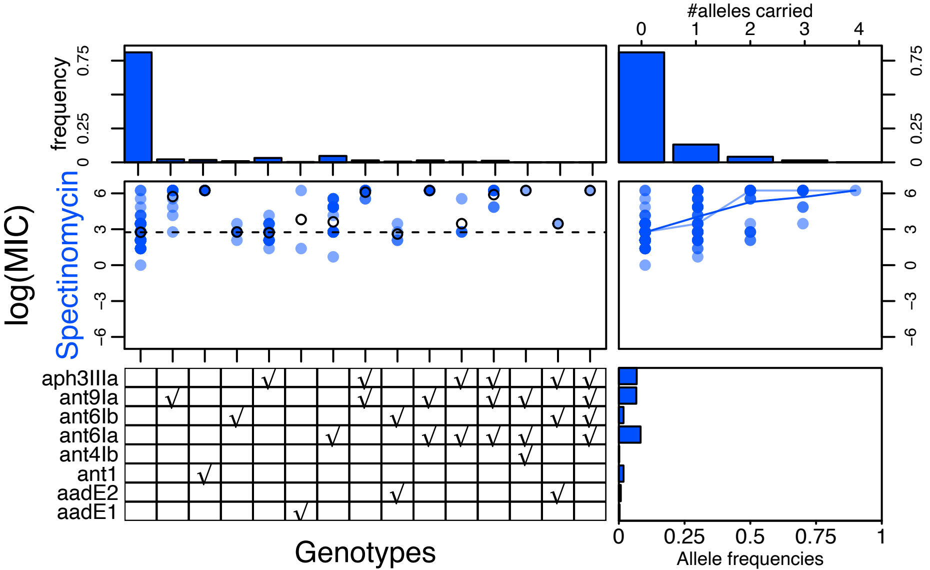
#alleles carried
0
1
2
3
4
frequency
0.75
0.25
0
log(MIC)
Spectinomycin
6
3
0
−3
−6
aph3IIIa
ant9Ia
ant6Ib
ant6Ia
ant4Ib
ant1
aadE2
aadE1
Genotypes
0
0.25
0.5
0.75
1
Allele frequencies

**Figure S6. The effects of candidate alleles on MIC for the aminoglycoside antibiotic, spectinomycin.** All details match Figure S2. As discussed in the main text, the candidate alleles *ant(6')-Ib* and *aph(3')-IIIa* seem to have little effect on MIC levels.

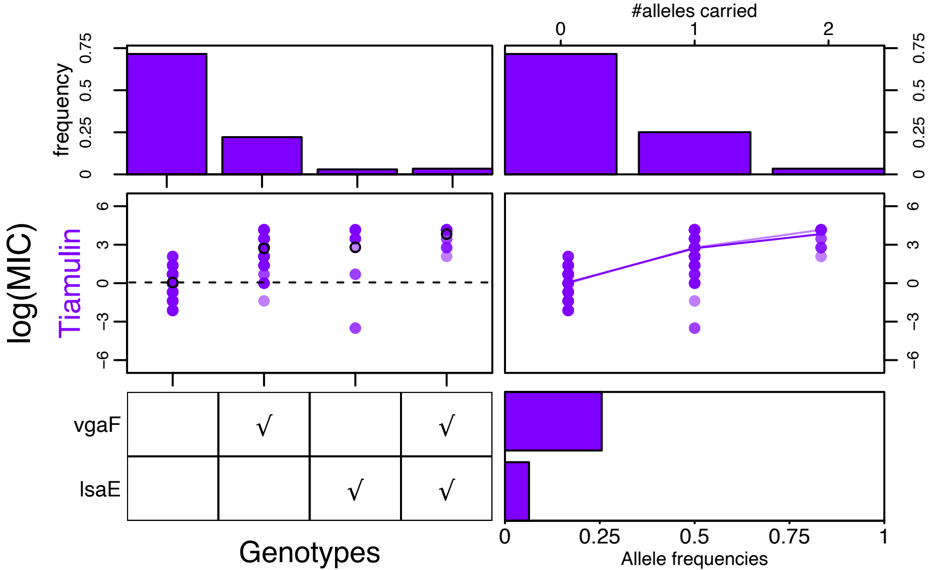

#alleles carried
0
1
2
frequency
0
0.25
0.5
0.75
log(MIC)
Tiamulin
6
3
0
−3
−6
vgaF
lsaE
√
√
√
√
0
0.25
0.5
0.75
1
Genotypes
Allele frequencies

**Figure S7. The effects of candidate alleles on MIC for the pleuromutilin antibiotic tiamulin.** All details match Figure S2.

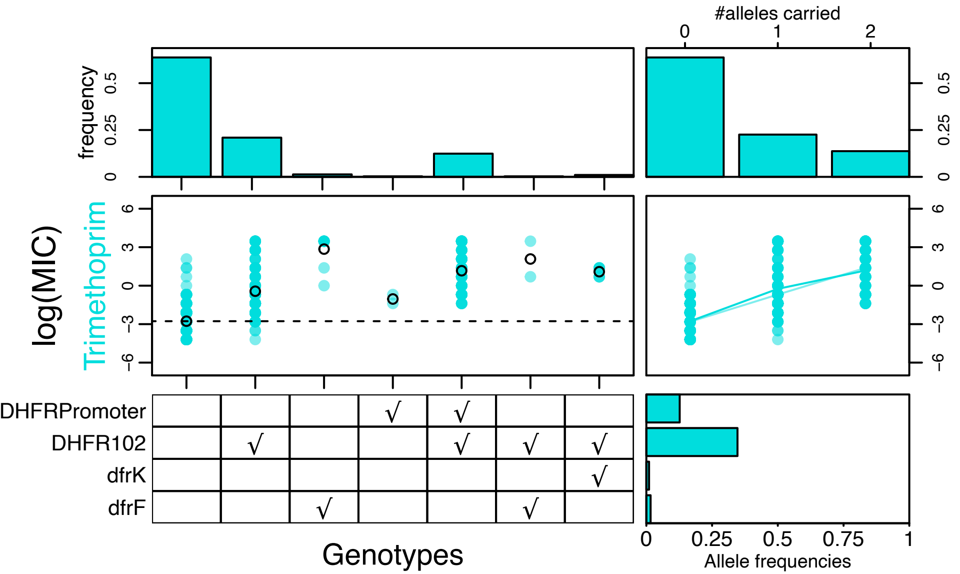
#alleles carried
frequency
log(MIC)
Trimethoprim
DHFRPromoter
DHFR102
dfrK
dfrF
Genotypes
Allele frequencies

**Figure S8. The effects of candidate alleles on MIC for trimethoprim (TMP).** All details match Figure S2.

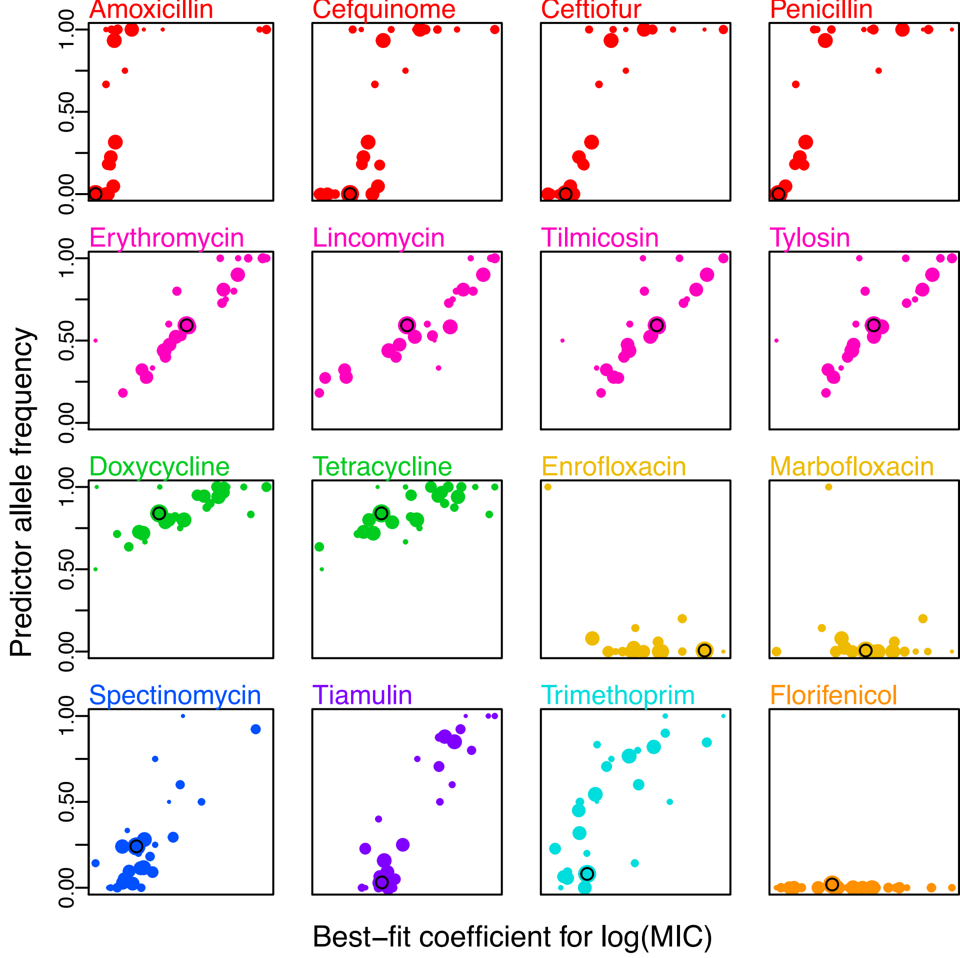

Amoxicillin
Cefquinome
Ceftiofur
Penicillin
Erythromycin
Lincomycin
Tilmicosin
Tylosin
Doxycycline
Tetracycline
Enrofloxacin
Marbofloxacin
Spectinomycin
Tiamulin
Trimethoprim
Florifenicol
0.00
0.50
1.00
Predictor allele frequency
Best–fit coefficient for log(MIC)

**Figure S9. Variation in the presence of candidate AMR determinants explains consistent differences between genetic clusters.** Each of 678 isolates was assigned to one of 30 genetic clusters (see methods), and linear models indicated significant differences in typical AMR levels between groups (Table 1). Each plot contains results for a single antibiotic, and compares the best-fit log(MIC) value for each genetic cluster, to the frequency of candidate AMR alleles for that antibiotic class, for isolates in that cluster. There is a clear tendency for clusters with higher MIC to have higher allele frequencies, except for the three antibiotics (enrofloxacin, marbofloxacin and florifenicol), where our candidate determinants have little predictive power, and our data set may represent a wild-type population. In each plot, point size corresponds to the number of isolates in each cluster, while a black circle indicates the BAPS4 cluster, which contains all of the isolates from Vietnam.

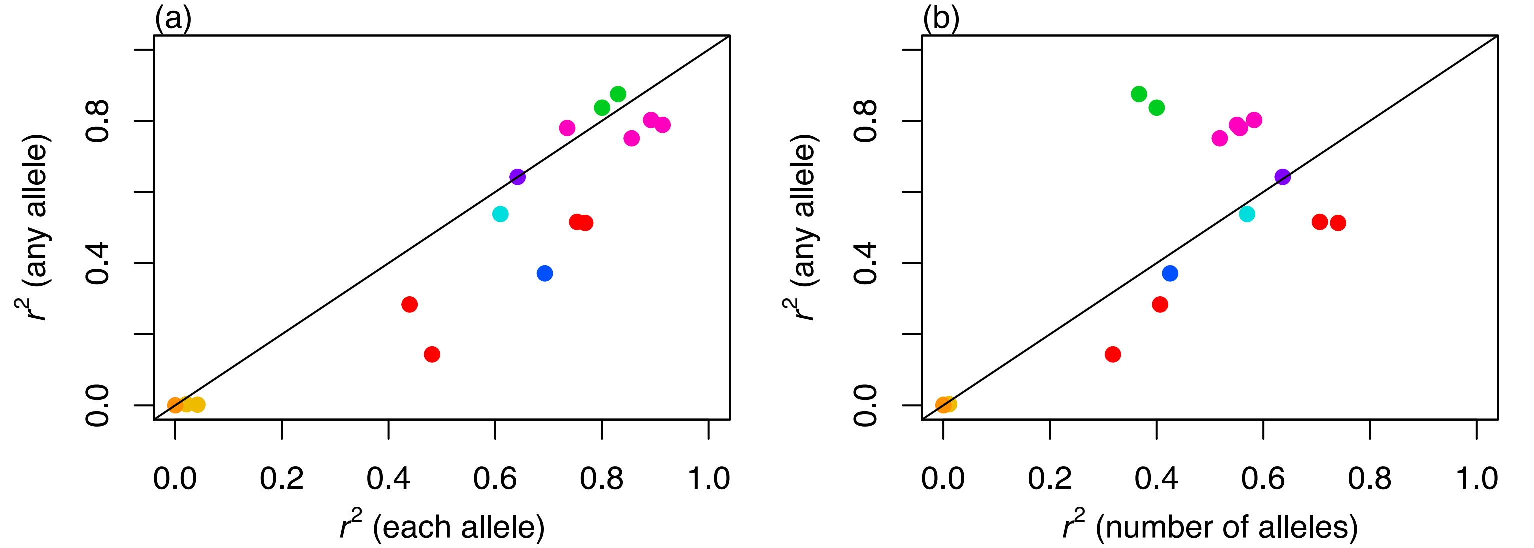
(a)
r^2 (any allele)
0.0
0.4
0.8
0.0
0.2
0.4
0.6
0.8
1.0
r^2 (each allele)
(b)
r^2 (any allele)
0.0
0.4
0.8
0.0
0.2
0.4
0.6
0.8
1.0
r^2 (number of alleles)

**Figure S10. Methods of using candidate determinants to predict MIC.** Each point compares the proportion of variance explained by a linear model including the candidate determinants for a given antimicrobial drug. For both panels, the value on the *y*-axis shows the $r^2$ for the simplest model, with single binary predictor variable, stating whether one or more candidate determinant is present or absent (these are the values shown in Figure 1). In the left-hand panel, these are compared to the $r^2$ values for a more complex model in which *each* of the candidate determinants is included as a separate binary predictor. This improved predictive power especially for the aminoglycoside spectinomycin (blue point), where some of our candidate predictors had no effect on MIC (see Figure S6 and main text); and for the $MLS_B$ class drugs (pink points), where some determinants worked only against macrolides or lincosamides (see Figure S3 and main text); and for the beta-lactams (red points) where alleles acted additively. The right-hand panel shows that results for this drug class could also be improved by predicting log MIC from the number of candidate determinants that were carried. By contrast, this strategy reduced predictive power for the $MLS_B$ class (pink) and tetracylines (green) where single determinants of large effect were sufficient to confer high MIC (see Figures S3 and S4).

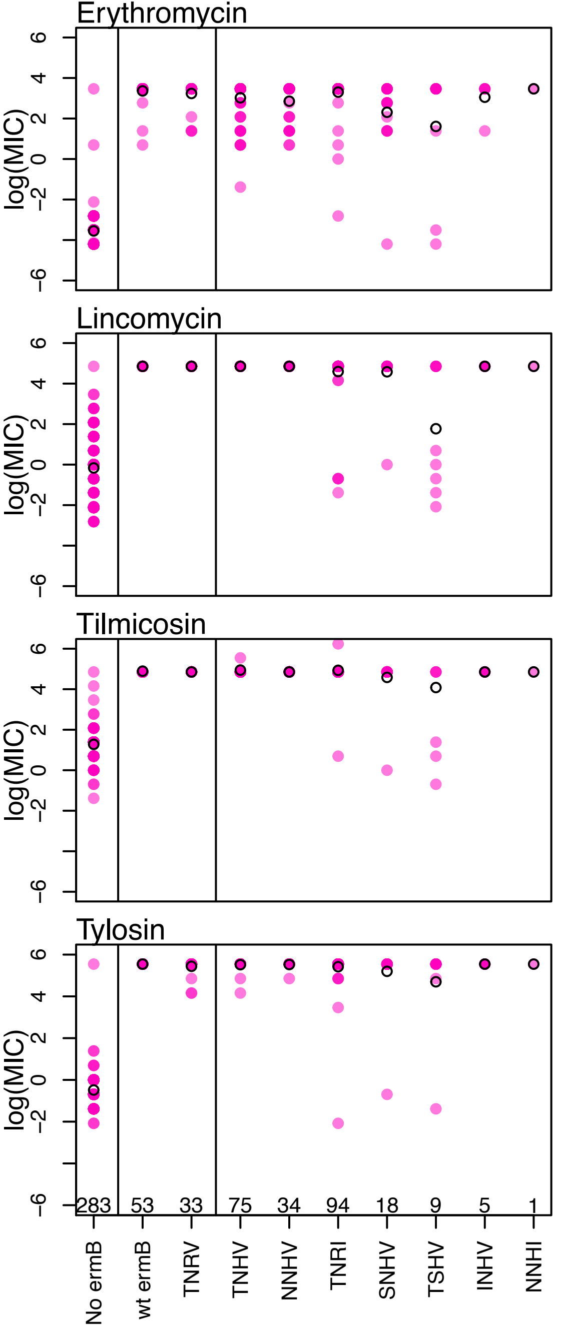
Erythromycin
Lincomycin
Tilmicosin
Tylosin
log(MIC)
6
4
2
0
−2
−6
283
53
33
75
34
94
18
9
5
1
No ermB
wt ermB
TNRV
TNHV
NNHV
TNRI
SNHV
TSHV
INHV
NNHI

**Figure S11. Allelic variation in *ermB* and unidentified sources of epistasis can affect MICs against macrolide-lincosamide-streptogramin B ($MLS_B$) antibiotics.** The first column ("No *ermB*") shows the log MIC values for the 283/678 isolates that contained neither *ermB* nor any other candidate determinants for $MLS_B$ drugs. The remaining columns show log MIC for some subset of the remaining 322 isolates that carried only *ermB* (i.e. no other candidate determinant for the $MLS_B$ class), and which had complete and successfully translatable sequences in our assemblies. *ermB* has 245 amino-acids, and 53/322 isolates carried the most common variant at all sites. We found that these wild-type *ermB* sequences ("wt *ermB*") had consistently high MICs. Rare amino-acid variants were segregating at 18/245 sites, but low MICs were found only in isolates that carried a rare variant at one or more of four positions: T75X, N100S, R118H and V226I. Accordingly, we classified each strain according the amino-acid state at these four positions. The 33/322 isolates that matched the wild-type at these positions ("TNRV") had high MIC. However, as shown by the remaining columns, there was no single *ermB* sequence that was consistently associated with low MIC. Altogether, then, the data suggest that MIC is affected both by allelic variation in *ermB* and by an epistatic factor elsewhere in the genome.

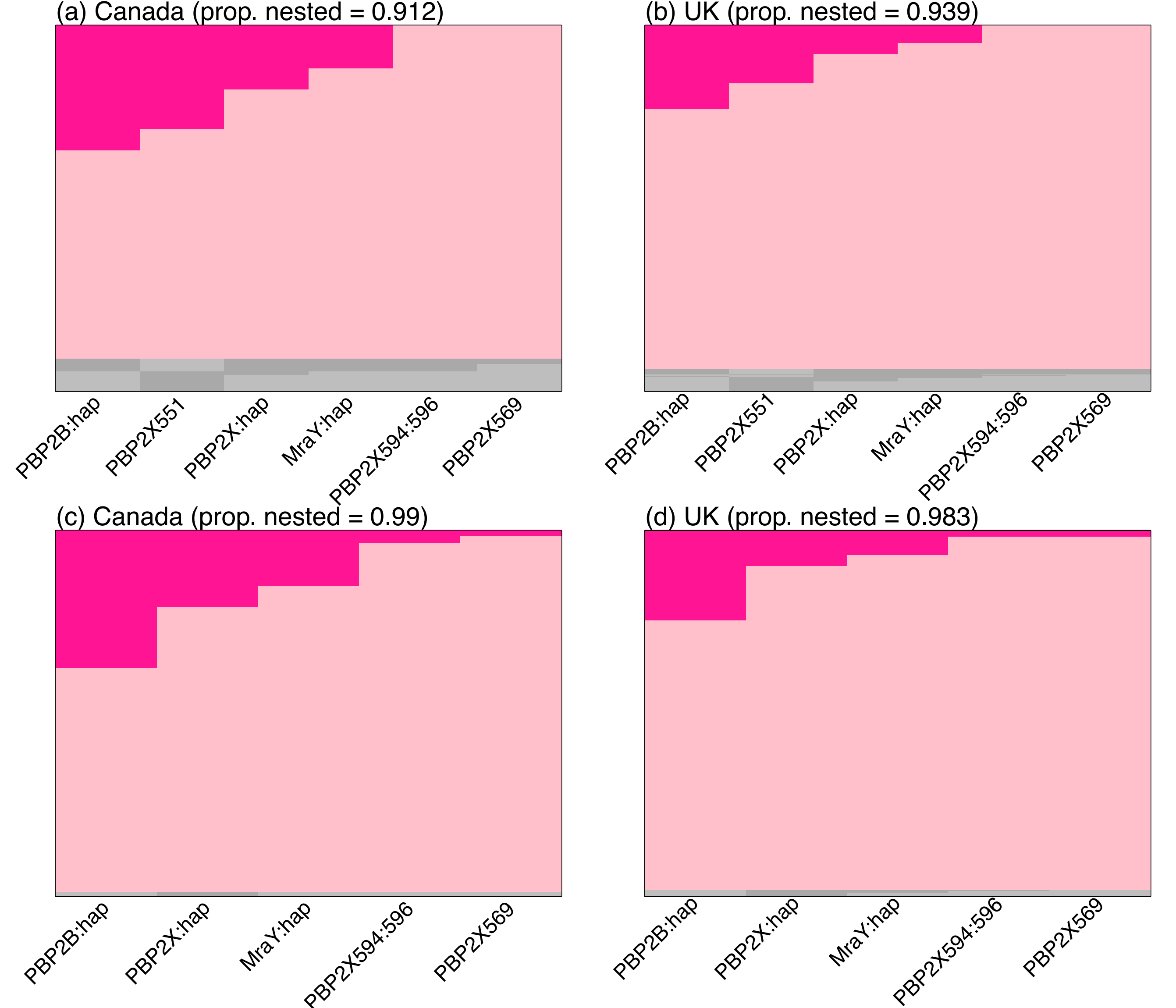
(a) Canada (prop. nested = 0.912)
PBP2B:hap
PBP2X551
PBP2X:hap
MraY:hap
PBP2X594:596
PBP2X569
(b) UK (prop. nested = 0.939)
PBP2B:hap
PBP2X551
PBP2X:hap
MraY:hap
PBP2X594:596
PBP2X569
(c) Canada (prop. nested = 0.99)
PBP2B:hap
PBP2X:hap
MraY:hap
PBP2X594:596
PBP2X569
(d) UK (prop. nested = 0.983)
PBP2B:hap
PBP2X:hap
MraY:hap
PBP2X594:596
PBP2X569

**Figure S12. High levels of “nestedness” suggest that resistance determinants to beta-lactams are acquired in a particular order.** All panels were generated using the methods and plotting conventions of [2] that were also used in the left-hand panel of Figure 4. The upper panels show that nestedness is high in the two largest subsets of our isolates, obtained from pigs from (a) Canada, or (b) the UK. The lower panels show that almost all isolates show a nested pattern when we exclude the (relative common) variant at site 551 in the PBP2X gene.

**Table S1 (separate data table). The 678 isolates in our collection with meta data, MIC values and candidate determinant presence/absence.**

| Variant | Antibiotic | MIC (mg/L) | Number of BAPs clusters | No of success | Exact Binomial one tailed *p* value | 95% CI |
|---|---|---|---|---|---|---|
| PBP2B_hap | Penicillin | ≥1 | 7 | 7 | 0.01563 | 0.59-1 |
| MraY_hap | Penicillin | ≥1 | 7 | 7 | 0.01563 | 0.59-1 |
| PBP2B_hap | Ceftiofur | ≥1 | 7 | 7 | 0.01563 | 0.59-1 |
| MraY_hap | Ceftiofur | ≥1 | 7 | 7 | 0.01563 | 0.59-1 |
| PBP2X_hap | Ceftiofur | ≥2 | 9 | 9 | 0.0039 | 0.66-1 |
| PBP2XT551S | Penicillin | ≥1 | 12 | 11 | 0.00634 | 0.066-1 |
| PBP2XT551S | Ceftiofur | ≥2 | 8 | 8 | 0.00781 | 0.063-1 |
| *vgaF* | Tiamulin | ≥8 | 13 | 13 | 0.00024 | 0.75-1 |
| *dhfr* promoter (A5G substitution and inserts within 1-30bp upstream) | TMP | ≥1 | 11 | 10 | 0.01172 | 0.58-0.99 |
| DHFRI102L | TMP | ≥0.25 | 20 | 19 | 4.01E-05 | 0.75-0.99 |

**Table S2. Binomial tests showing that novel AMR variants are independently associated with MIC in different genetic clusters.**

**Table S3 (separate data table). The 401 additional isolates used to estimate population structure with their genetic BAPs cluster and their data availability.**

### References


1. Murray GGR, Charlesworth J, Miller EL, Casey MJ, Lloyd CT, Gottschalk M, Tucker AW, Welch JJ, Weinert LA: **Genome Reduction Is Associated with Bacterial Pathogenicity across Different Scales of Temporal and Ecological Divergence**. *Molecular biology and evolution* 2021, **38**(4):1570-1579.
2. Lehtinen S, Blanquart F, Lipsitch M, Fraser C, with the Maela Pneumococcal C: **On the evolutionary ecology of multidrug resistance in bacteria**. *PLoS pathogens* 2019, **15**(5):e1007763.
